# Centromere-associated retroelement evolution in *Drosophila melanogaster* reveals an underlying conflict

**DOI:** 10.1101/2022.11.25.518008

**Authors:** Lucas W. Hemmer, Sherif Negm, Xuewen Geng, Cécile Courret, Beatriz Navarro-Domínguez, Iain Speece, Xiaolu Wei, Eddyson Altidor, James Chaffer, John S. Sproul, Amanda M. Larracuente

## Abstract

Centromeres are chromosomal regions essential for coordinating chromosome segregation during cell division. While centromeres are defined by the presence of a centromere-specific histone H3 variant rather than a particular DNA sequence, they are typically embedded in repeat-dense chromosomal genome regions. In many species, centromeres are associated with transposable elements, but it is unclear if these elements are selfish or if they play a role in centromere specification or function. Here we use *Drosophila melanogaster* as a model to understand the evolution of centromere-associated transposable elements. *G2/Jockey-3* is a non-LTR retroelement in the *Jockey* clade and the only sequence shared by all centromeres. We study the evolution of *G2/Jockey-3* using short and long read population genomic data to infer insertion polymorphisms across the genome. We combine estimates of the age, frequency, and location of insertions to infer the evolutionary processes shaping *G2/Jockey-3* and its association with the centromeres. We find that *G2/Jockey-3* is an active retroelement targeted by the piRNA pathway that is enriched in centromeres at least in part due to an insertion bias. We do not detect signatures of positive selection on any *G2/Jockey-3* insertions that would suggest than individual copies are favored by natural selection. Instead, we infer that most insertions are neutral or weakly deleterious both inside and outside of the centromeres. Therefore, *G2/Jockey-3* evolution is consistent with it being a selfish genetic element that targets centromeres. We propose that targeting centromeres helps active retroelements escape host defenses, as the unique centromeric chromatin may prevent targeting by the host silencing machinery. At the same time, centromeric TEs insertions may be tolerated or even beneficial if they also contribute to the transcriptional and chromatin environment. Thus, we suspect centromere-associated retroelements like *G2/Jockey-3* reflect a balance between conflict and cooperation at the centromeres.

## INTRODUCTION

Centromeres are important chromosomal structures for kinetochore assembly and proper chromosome segregation during cell division. While centromeres are usually in highly repetitive regions, the role of DNA sequences in centromere function or establishment remains unclear (reviewed in [1]). Across many organisms, centromeres are associated with satellite DNAs (satDNAs), large blocks of tandemly repeated DNA sequences (reviewed in [2, 3]). Centromere-associated tandem repeats are capable of initiating centromere assembly in fission yeast [4] and humans [5]. However, centromeres are not defined by any particular DNA sequences. They are instead defined epigenetically by CENP-A, a specific variant of histone H3 [6–11]. Centromere domains consist of unique chromatin where H3 nucleosomes are interspersed with CENP-A nucleosomes [12–15]. Repeat sequences are not always required for centromere formation, as neocentromeres can form outside of centromeric DNA in the absence of satDNA sequences [16–18] and some species have centromeres devoid of repeats (*e.g.* satellite-free centromeres in the Equus genus [19]). Neocentromeres have the potential to replace older centromeres with implications for karyotype evolution and speciation [20–22].

Centromeres and pericentromeric heterochromatin are also enriched in transposable elements (TEs)— selfish genetic elements that mobilize in genomes. Evidence for specific TE-centromere associations has emerged in the last two decades, including in grasses [23, 24], rice [25, 26], maize [27], marsupials [28, 29], bats [30], gibbons [31, 32], and humans [18, 33, 34]. However, it is unclear how, or if, TEs contribute to centromere biology for most species [35]. Centromeres may offer a ‘safe haven’ to TEs that jump there and for the genome [36, 37], as centromere insertions are unlikely to disrupt centromere or gene function [35]. At the same time, centromeric TEs may contribute to important aspects of centromere evolution and biology. Homology between some TEs and centromeric tandem repeats [3, 38] suggests that TEs can contribute to centromere evolution by giving rise to new centromeric satDNAs [39]. TEs may also play roles in centromere function. These roles may be indirect, such as maintaining the constitutive pericentromeric heterochromatin, which is important for centromere function [40–42]. However, more direct roles are also possible. Some transcription of centromeric regions is necessary for recruitment of CENP-A and establishing centromere location [43–47]. Centromeric transcripts may also facilitate recruitment of kinetochores to centromeric DNA [48, 49]. Given that TEs contain sequences important for promoting transcription [50], their presence in centromeres may contribute to centromere specification and function [35]. In most cases it is unclear whether TEs are a part of the CENP-A domain corresponding to functional centromeres or just associated with adjacent, repeat-rich regions of chromosomes where functional centromeres are often embedded. Advances in single molecule long-read sequencing technology now allows for detailed study of centromere organization at the sequence level to provide insights into centromere structure, composition, and evolution [51–56].

With its compact genome and expansive genetic toolkit, *Drosophila melanogaster* is an excellent model organism to study centromere evolution [57]. A recent study of centromere organization in *D. melanogaster* revealed that CENP-A is bound to TE-rich islands of complex DNA embedded within large arrays of satellite repeats [53]. The composition of each centromere island is unique. The only sequence element shared among all centromeres is a single non-LTR retroelement in the *Jockey* superfamily, *G2/Jockey-3.* However, *G2/Jockey-3* is not exclusively located in centromeres; about 40% of all insertions are found outside of the centromere islands [53]. Polymorphic *G2/Jockey-3* insertions [58] and transcripts from published RNAseq datasets [59–61] suggest this element may still be active in *D. melanogaster* [53]. *G2/Jockey-3* is also found at the centromeres in *D. simulans*, a close relative of *D. melanogaster*, suggesting that its centromere association may be broadly conserved [53].

It is unclear whether *G2/Jockey-3* is an important centromere component in *D. melanogaster* or if they are instead purely selfish TEs. If *G2/Jockey-3* are important for centromere specification or function, we may detect evidence for positive natural selection on centromeric copies. Alternatively, *G2/Jockey-3* may be a selfish element that targets insertions in centromeres but are otherwise selectively neutral or weakly deleterious. These hypotheses are not necessarily mutually exclusive. Here we investigated the age and frequency of *G2/Jockey-3* insertions across *D. melanogaster* genomes to determine if its evolution is consistent with a role in centromere function or underlying selfish behavior. Our analyses suggest that *G2/Jockey-3* insertions are largely neutral or weakly deleterious both inside and outside centromeres, suggesting that they are selfish genetic elements. Our results also suggest that *G2/Jockey-3* has had a weak insertion bias toward centromeres. Although there is no evidence for positive selection on individual *G2/Jockey-3* centromeric insertions, insertion of active TEs like *G2/Jockey-3* may still contribute to transcription and the unique chromatin environment at centromeres. We propose that there may be a balance between conflict and cooperation between active TEs like *G2/Jockey-3* and centromeres.

## RESULTS

### 1. A new consensus sequence for G2/Jockey-3

Prior to analyzing the distribution of *G2/Jockey-3* insertion polymorphisms in *D. melanogaster* populations, we first confirmed the consensus annotation. We analyzed all *G2/Jockey-3* insertions in our *D. melanogaster* genome assembly [53] and found two lines of evidence that the annotation needed updating: 1) 25 of the 167 *G2/Jockey-3* sequences in the genome shared up to 92 bp of upstream sequence at the 5’-end of the annotation; and 2) our reanalysis of PROseq data [62] suggests that *G2/Jockey-3* transcription starts 92 bp upstream of the published start site (Figure S1A). This upstream region is also enriched for GO terms for RNA polymerase II transcription factors and other transcription factor activity (Figure S1B). We therefore extended the original *G2/Jockey-3* consensus sequence by 92 bases (File S1).

*G2/Jockey-3* is a part of the *Jockey* superfamily of non-LTR retroelements, which typically contain two open reading frames (ORFs) (Figure S1A). While ORF1 is generally less conserved [63], ORF2 is well-conserved across *Jockey* elements and encodes for a reverse transcriptase (RT) and an endonuclease. NCBI ORF finder detected both ORFs within *G2/Jockey-3*. ORF1 is predicted to produce a 427 amino-acid protein approximately 186 bp downstream of the 5’ end. The NCBI Conserved Domains identified a PRE_C2GC super family (cl11655, E-value 9.22E-18) within ORF1 and while the function is unknown, it is associated with Cys2His2 type zinc fingers found in putative GAG proteins in other arthropod species. The predicted ORF2 overlaps with the last four bp of ORF1 and contains a domain corresponding to endonuclease (EEP Superfamily Accession cl00490, E-value 7.21E-18) and RT (RT_like Superfamily Accession cd01650, E-value 4.62E-56). *G2/Jockey-3* ends in a poly(A) signal.

### 2. Overview of data sets analyzed

We studied the evolutionary dynamics of *G2/Jockey-3* and other TEs by investigating the patterns of insertion polymorphisms both in a single panmictic population, and in individuals from a wide geographic range (Figure 1, Table S1). We conducted a survey of TE dynamics and genetic variation in 135 samples from the *Drosophila melanogaster* Genetic Reference Panel (DGRP) collected from a single population in North Carolina with whole genome Illumina sequence data [64, 65] and a subset of long read data from the same population for validation [66] (see Supplemental Text). For the samples from a wide geographic range, we used both long- and short-read data from 13 founder lines of the *Drosophila* Synthetic Population Resource (DSPR) and Oregon-R [67]. We also analyzed long-read PacBio data from five Global Diversity Lines (GDL) samples from different continents [68]. We generated *de novo* genome assemblies of each line to determine the organization of the centromere islands originally described in the reference genome for comparison (Table S2) and predicted insertions in short read data with the McClintock pipeline [69]. We assessed the reliability of our TE insertion predictions by looking for concordance between short and long read predictions (see Supplemental Text) and through PCR validation of a subset of insertions. We confirmed 64% (67 / 105) of the *G2/Jockey-3* insertions detected by McClintock are found in the long-read samples we examined (File S2). PCR confirmed 80% (28 / 35) of *G2/Jockey-3* insertions predicted in the GDL lines, and ~67% (12 / 18, File S3) of the centromere insertions. Based on our assessment, we limit our analysis to short read datasets with >20X centromere coverage (Table S3) and TEs predicted by more than one TE caller implemented in McClintock (Supplemental Text).

**Figure 1:**
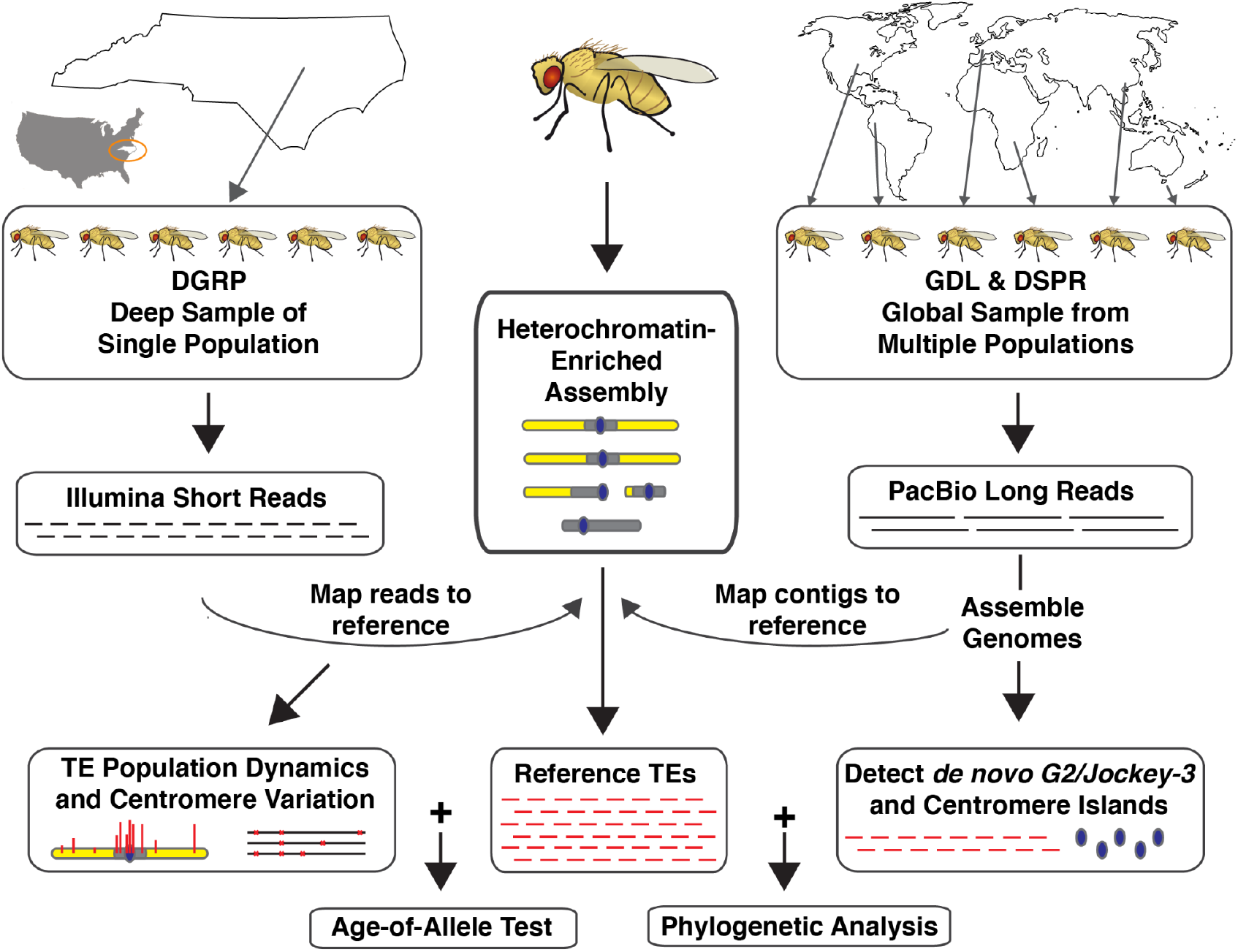
The summary of data and analyses using one source or a combination of sources in this paper. A heterochromatin-enriched assembly was originally used to discover centromere islands in Chang et al. [53] and identify reference TE insertions. We used short reads from the DGRP to identify reference and *de novo* insertions to study TE dynamics within a single population from North Carolina (map source: R package usmaps https://cran.r-project.org/web/packages/usmap/). We generated genome assemblies for 18 samples from global populations to uncover the centromere island sequences and *de novo* elements (map source: www.outline-world-map.com). We combined the reference sequences with population frequencies to infer selection with an age-of-allele test [70]. We also combined reference TE sequences with *de novo* elements uncovered in our assemblies for phylogenetic analysis. The fly illustrations are modified images from Wikipedia Commons (https://commons.wikimedia.org/wiki/File:Drosophila-drawing.svg).

### 3. G2/Jockey-3 is active in Drosophila melanogaster populations

Previous studies of TE polymorphism in *D. melanogaster* are consistent with *G2/Jockey-3* (previously annotated as *G2*) being recently active [58, 71, 72]. To determine if *G2/Jockey-3* is active in the DGRP, we detected TE insertion polymorphisms with the McClintock pipeline [69] and classified each insertion based on their presence (a “reference” insertion) or absence (a non-reference or *“de novo*” insertion, Figure 2) in our reference genome [53, 73]. *G2/Jockey-3* showed approximately five *de novo* insertions per line (File S4-S5), which is considerably fewer than the most highly active *Drosophila* TEs. For example, we estimate *Hobo* and *Jockey-1* have approximately 35 *de novo* insertions per line (consistent with [58]). However, the fraction of *de novo* insertions as percentage of all insertions (39% or 733 / 1124 for *G2/Jockey3*) is comparable to other highly active elements [72] such as *Doc* (25%; 2944 / 3826), *Jockey* (24%; 1308 / 5448), *I* element (27%; 658 / 2364), and *Copia* (88%; 1083 / 1226) (File S4).

**Figure 2:**
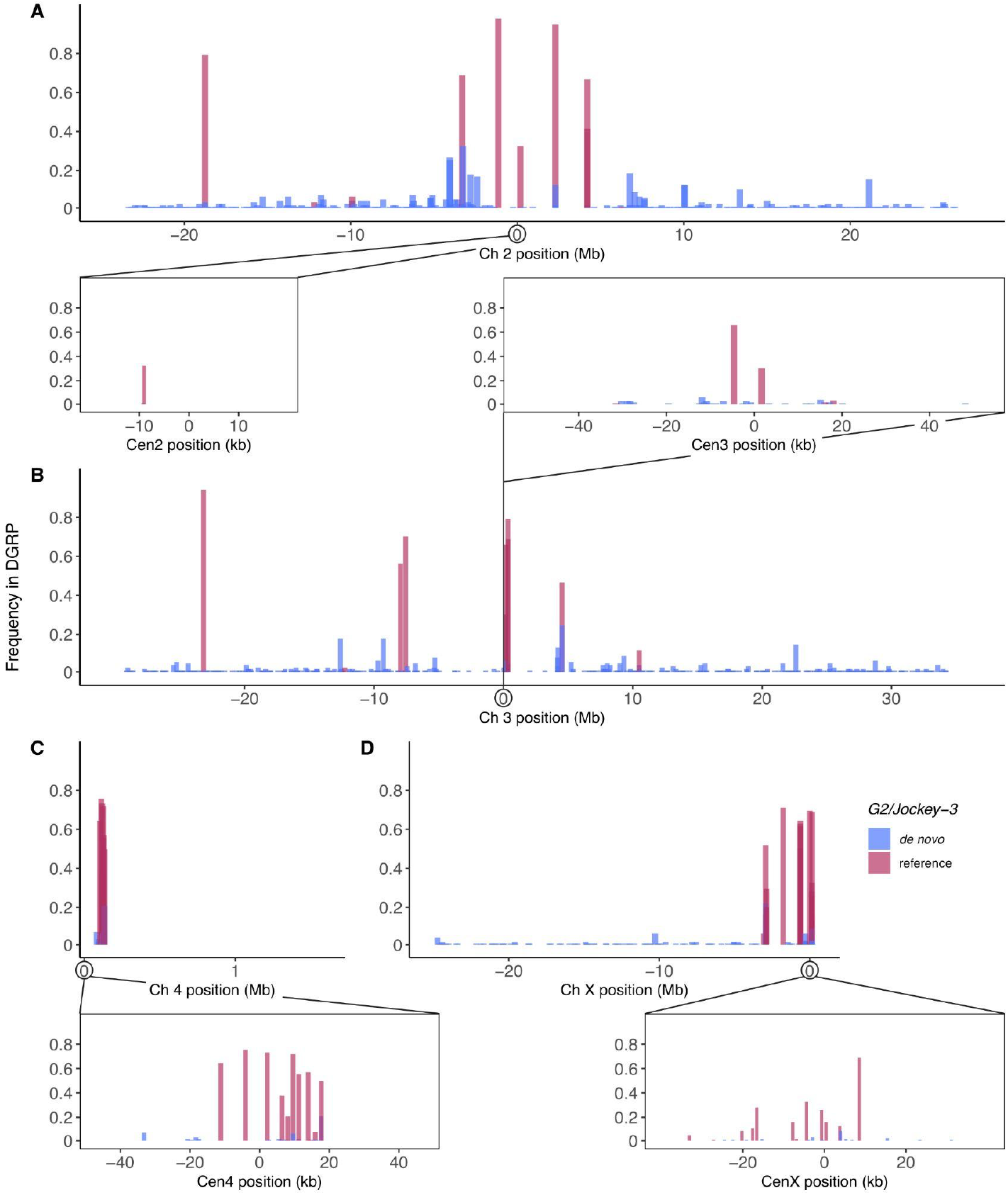
The distribution and frequency of *G2/Jockey-3* insertions in *D. melanogaster.* The height of the bars corresponds to frequency of an insertion detected in the DGRP samples and the color denotes whether an insertion is found in the reference genome or is *de novo.* Each A-D subpanels depict a chromosome (Ch 2, Ch 3, Ch 4, Ch X) from the heterochromatin-enriched assembly of *D. melanogaster* genome [53, 73], with a window zoomed in on the corresponding centromere (*e.g.*, X chromosome centromere = CenX, *etc*).

We detected insertions in the centromeres, here defined as the centromere islands identified in Chang et al. [53]. Of the 18 different TE families detected within centromeres, *G2/Jockey-3* is the most common. Within centromeres, *G2/Jockey-3* has the highest percentage of *de novo* (32%; 15/47) and reference (72%; 28 / 39, Figure 2) TE insertions. Only *G2/Jockey-3* and *Doc* have more than 10 unique insertions (File S4). Other *de novo* centromere insertions include active families such as the *F* element, *R1* and *Roo.*

Many active TEs are targeted by small, 24-30-nt long Piwi-interacting RNAs (piRNA) in the germline [74, 75]. These piRNAs target TEs for both transcriptional and post-transcriptional silencing. In the *D. melanogaster* germline, piRNAs are produced from precursor transcripts that originate from discrete loci in the genome that contain clusters of nested TEs [76] called piRNA clusters [77]. Subsequent amplification of the piRNA signal comes from the piRNA-guided cleavage of active TEs through the ‘ping-pong’ cycle [77, 78] generating a signature of 10-nt overlap between sense and antisense piRNAs. Consistent with *G2/Jockey-3* being a recently active element, we detect insertions of *G2/Jockey-3* in the *38C1* (2 copies) and *38C2* (9 copies) piRNA clusters, and *G2/Jockey-3*-derived piRNAs in the germline. *G2/Jockey-3* is ranked in the top 4% and top 18% of TE-derived piRNAs in testes and ovaries, respectively (Table S4). The resulting *G2/Jockey-3* piRNAs show signatures of ping-pong in ovaries (Z-score = 7.11; data from [60, 79, 80]) but not in testes (Z-score = 0.15; data from [81–83]), consistent with other testis piRNAs. Taken together, these results suggest that *G2/Jockey-3* is an active TE enriched in centromere islands.

### 4. Young G2/Jockey3 insertions are enriched in centromeres

To understand if the observed enrichment of *G2/Jockey3* at centromeres is due to insertion bias, we studied insertion frequencies across the genome. We reasoned that an enrichment of putatively young *G2/Jockey-3* at centromeres may reflect an insertion preference. Young, recently inserted TEs are expected to occur at lower frequencies than older TEs. We thus examined the distribution of rare insertions (<10% population frequency) of *G2/Jockey-3* and other TE families in the DGRP. We found that rare *G2/Jockey-3* insertions are significantly enriched in centromeres compared to other regions relative to their size (19 centromere insertions; Permutation tests: n_perm_ 10,000, *P* < 0.0001; *Z*-score = 26.736 for whole genome; *P* < 0.0001; *Z*-score = 23.895, excluding heterochromatin windows; *P* < 0.0001; *Z*-score = 17.354, excluding euchromatin windows; Figure 3). *R1* elements are the only other TE enriched in centromeres (4 centromere insertions; Permutation tests: n_perm_ = 10,000, *P* < 0.0001; *Z*-score = 6.892 for whole genome; *P* < 0.0001; *Z*-score = 5.92, excluding heterochromatin windows*; P* = 0.0187; *Z*-score = 2.605, excluding euchromatin windows; Figure 3), but to a lesser degree than *G2/Jockey-3*. Among rare insertions, *G2/Jockey-3* is by far the most enriched TE in centromeres (39% of all rare insertions).

**Figure 3:**
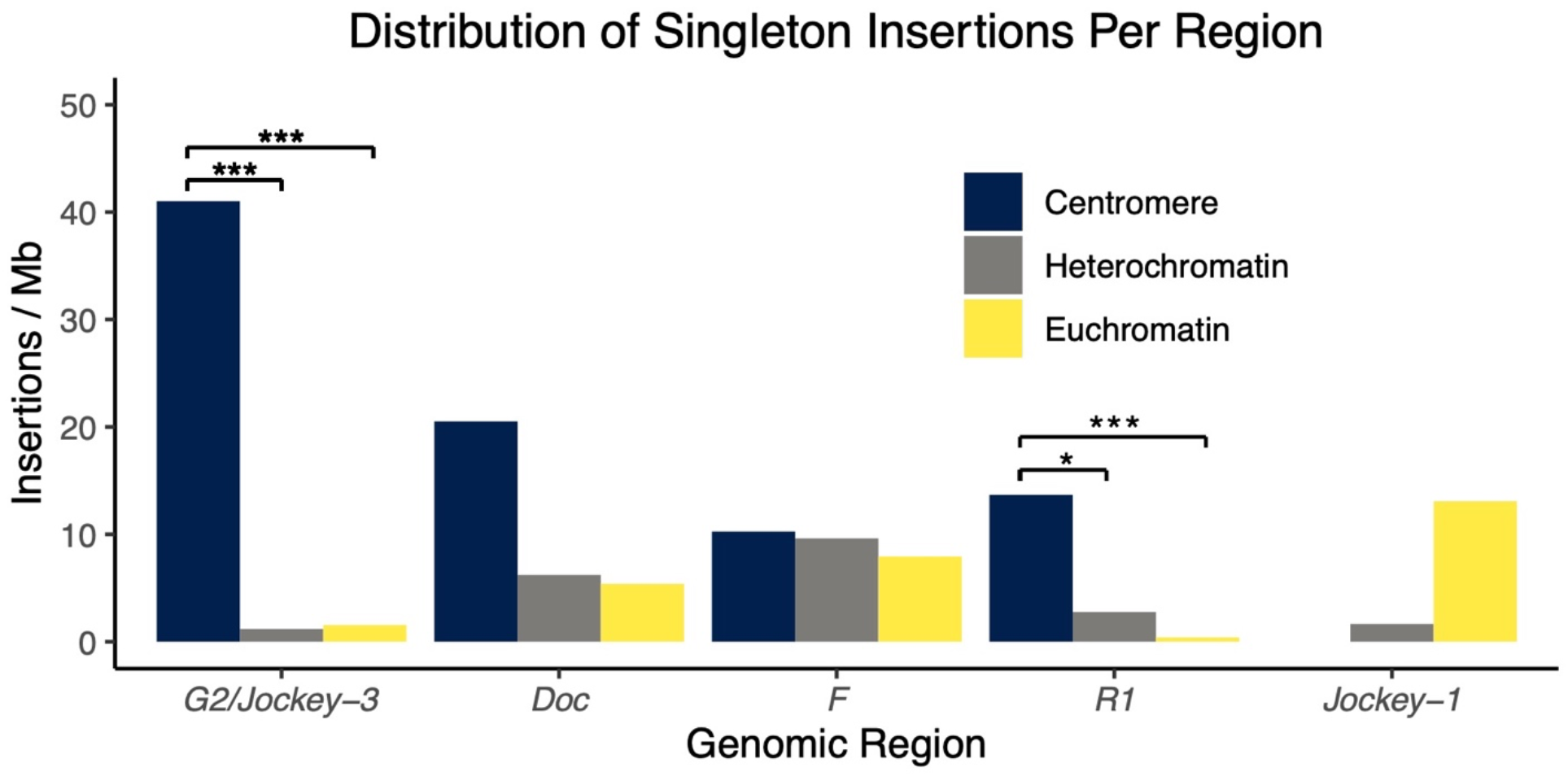
Young *G2/Jockey-3* insertions are highly enriched in centromeres relative to heterochromatin and euchromatin. The distribution of rare insertions (< 10% of the population) per Mb between the centromere islands, heterochromatin, and euchromatin regions for several TE families in DGRP lines. Significant differences are noted as: * *P* < 0.05, *** *P* < 0.0001 in the permutation test.

We confirmed these patterns using an orthogonal approach with long-read genome assemblies of flies from a wide geographic distribution (*i.e.*, founder DSPR lines, GDL lines, and our reference genome; Table S1). Here we identified putatively young *G2/Jockey-3* insertions based on sequence divergence instead of frequency. We classified young insertions as those with < 1% divergence from the consensus sequence, or with < 2% divergence from the consensus and found within a single genome assembly. Insertions classified as young were found in less than 50% of the assemblies, had fewer unique substitutions, and were more likely to have a complete ORF than older fragments (Figure 4, FET *P* = 3.93E-12, File S6, File S7). Consistent with our short-read DGRP findings (Figure 3), we found significant relative enrichment in centromeres (Permutation tests n_perm_ = 10,000: *P* < 0.0001; *Z*-score = 15.633 for whole genome; *P* < 0.0001; *Z*-score = 7.505 excluding euchromatin windows), even with some of the centromere island sequences missing in the long-read assemblies (Table S2).

**Figure 4:**
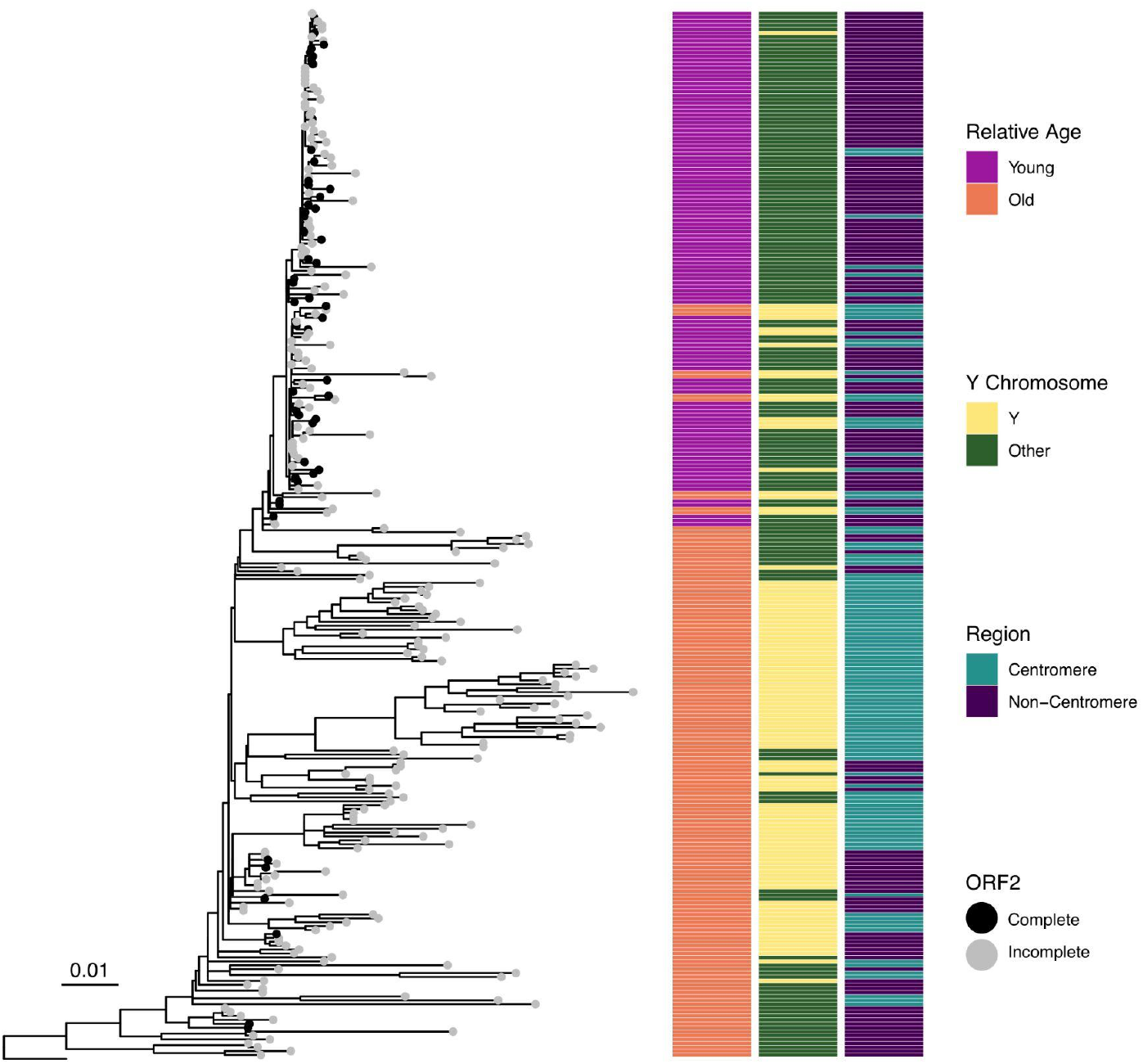
Young *G2/Jockey-3* insertions share a common origin. A maximum likelihood phylogenetic tree with all non-redundant *G2/Jockey-3* sequences larger than 10% of the full-length (400 bp) in the reference genome [53] and long-read assemblies of founder DSPR and GDL lines, with the *D. simulans* consensus as the outgroup. Black circles denote “complete” *G2/Jockey-3* elements containing a full ORF2; gray circles are incomplete elements. The columns and colors correspond to characteristics of each insertion including chromatin region (centromere or not), relative age, and presence on Y chromosome. Young insertions, determined using an orthogonal approach to the Figure 3 analysis, are those with < 1% divergence from the consensus sequence or < 2% and found within a single genome assembly.

These results suggest that *G2/Jockey-3* has had at least some insertion preference for centromeres. As insertion bias can be associated with a target site sequence, we searched for motif enrichment at insertions sites. Although we found an enrichment of AT-rich simple motifs like “TAGTTTT” and rDNA-associated sequences (*e.g.* 18S and non-transcribed spacers; Figure S2) within 100 bases of predicted insertion sites, these sequences are generally abundant in repeat-dense regions of the genome and not specifically enriched at *G2/Jockey-3* insertions (File S6, File S8; MEME *E*-value = 3.8E-16).

### 5. Skew toward high frequency copies of G2/Jockey-3 in the centromere

To test if *G2/Jockey3* insertions are retained at centromeres by natural selection, we studied their frequencies. Because most TE insertions are deleterious in the euchromatin, they are found at low population frequencies [84]. In contrast, TEs in heterochromatin are allowed to reach higher population frequencies because recombination is infrequent, and natural selection is less efficacious (reviewed in [85–87]). To infer differences in the strength and direction of natural selection in different genomic regions, we examined the frequency spectra of a subset of TE families in centromeres, heterochromatin, and euchromatin (Figure 5, Figure S3). If natural selection favors centromeric insertions, we should observe a skew toward high frequency variants in centromeres compared to heterochromatin. We compared the frequency spectra of two TE families which had five or more insertions in the centromeres: *G2/Jockey-3* and *Doc.* As expected, both elements showed an enrichment for rare insertions in the euchromatin, consistent with TEs being generally deleterious in gene-rich regions (Figure 5; FET *P* < 0.001). Similarly, we observed a slight skew toward higher frequency *G2/Jockey-3* and *Doc* insertions (*e.g.* common/intermediate) in heterochromatin and centromeres compared to euchromatin (Figure 5, FET *P* < 0.001), suggesting relaxed constraints or a reduced efficacy of natural selection outside of euchromatic regions. Other TE families found in centromeres are also enriched for low frequency insertions in the euchromatin (Figure S3). However, we did not observe differences between the frequency spectra of the euchromatin and centromeres, suggesting most TEs may be deleterious in these regions (Figure S3).

**Figure 5:**
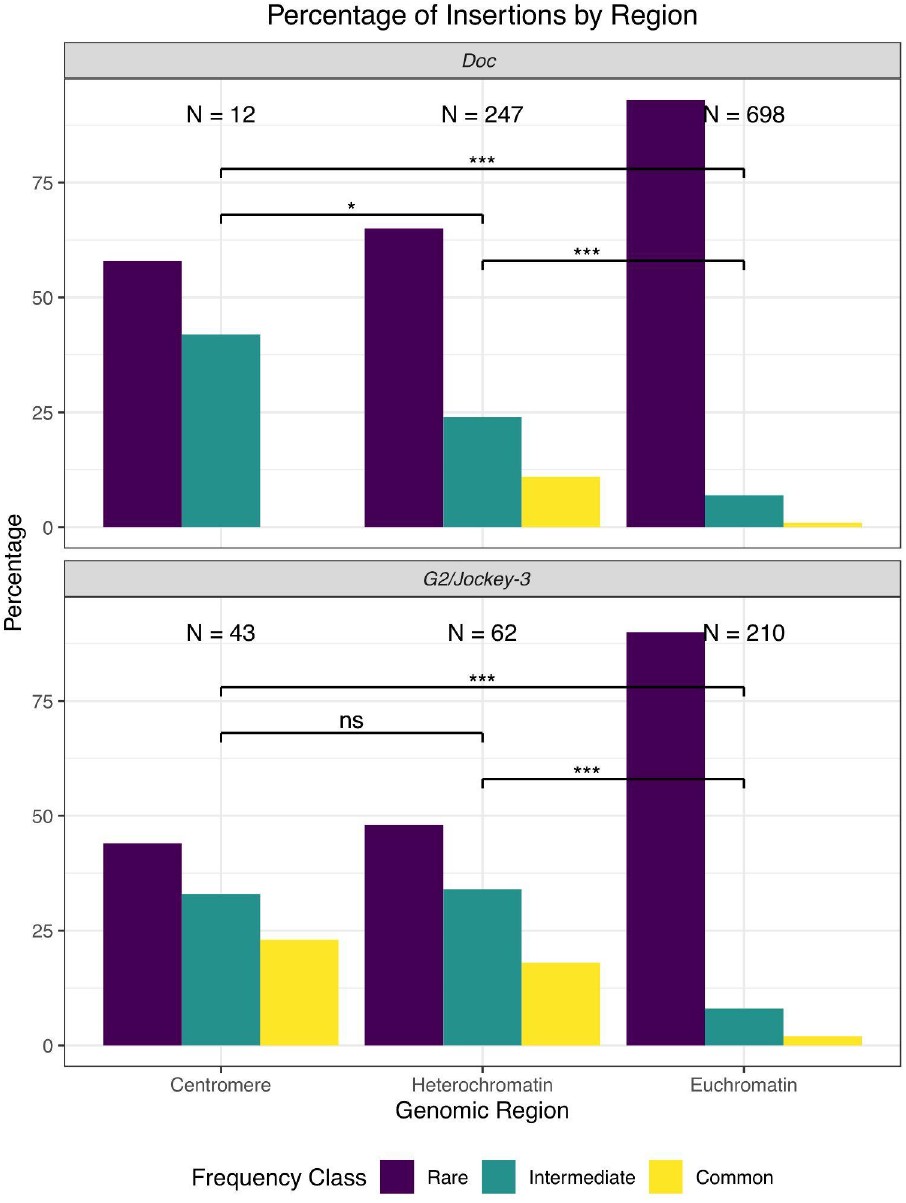
*G2/Jockey-3* has higher frequency variants in centromeres. TE frequencies were binned into rare (less than 10% of all samples), intermediate (10-50% of all samples), and common (> 50% of all samples). The frequency distributions of the four TE families with > 10 unique insertions in the centromeres were compared among the centromeres, heterochromatin, and euchromatin. Fisher’s Exact Tests between regions, Bonferroni adjusted two-tailed P-value * *P* < 0.05; *** *P* < 0.001; ns is not significant.

The frequency spectra of *G2/Jockey3* does not appear significantly different between centromeres and heterochromatin. However, *G2/Jockey-3* has more “common” insertions in centromeres relative to other TEs, including *Doc* (Figure 5; Figure S3). This could be due to relaxed constraints allowing *G2/Jockey-3* insertions to rise to higher frequencies over time or positive selection for some *G2/Jockey-3* insertions in centromeres.

### 6. G2/Jockey-3 ages suggest a historical burst of activity

Centromeric *G2/Jockey-3* copies are generally older—within a range of 2-7% divergence from the consensus—than those in heterochromatin (MWU, *P* < 0.001, Figure 6A), but not significantly different than euchromatic copies (MWU, *P* > 0.5; Figure 6A). This pattern suggests that most centromeric copies of *G2/Jockey-3* resulted from a historical burst of activity. This burst of activity is consistent with general patterns observed at other non-LTR retroelements [72, 88]. Other TE families show a similar age distribution between heterochromatin and the centromeres (Figure S4). These copies are also older and highly fragmented. In general, both the lack of recombination and lack of strong purifying selection against TE insertions in centromeres and heterochromatin may allow these insertions to persist over longer evolutionary time periods [89].

**Figure 6:**
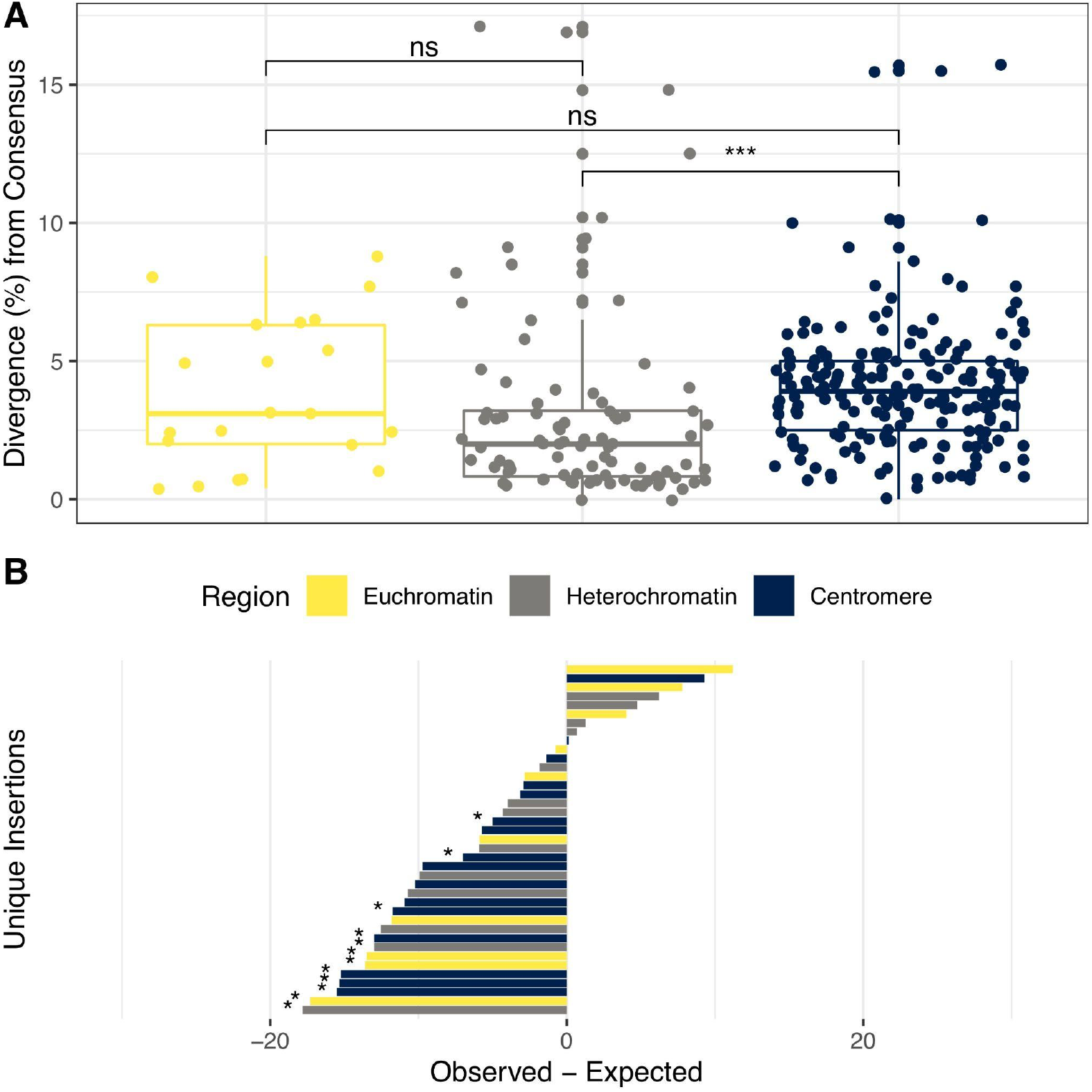
*G2/Jockey-3* copies are either as frequent or less frequent than expected given their age. A) The number of *G2/Jockey-3* insertions grouped by divergence from the consensus sequence split between the euchromatin, heterochromatin, and centromeres. Two-tailed Mann Whitney U tests *** *P* < 0.001, ns is not significant. B) The difference between observed and expected frequency of *G2/Jockey-3* copies in the DGRP. The *P*-values are calculated as the sum of the probabilities that an insertion is less frequent than expected (if observed - expected < 0) or more frequent than expected (observed - expected > 0). * *P* < 0.1, a stringent threshold given the reduced power to detect deviations from neutrality as stated in Blumenstiel et al. [70].

We can combine information about the TE’s age and frequency to infer if, and how, its history is consistent with natural selection. In the absence of selection, the frequency of a TE insertion should correlate with its age: *e.g.*, older TEs have more time to drift to higher population frequencies than young TEs and thus accumulate more unique mutations. Natural selection can cause deviations from expected frequencies based on the age. While negative selection might cause TEs to have lower population frequency than expected given their age, positive selection might increase their frequencies. We therefore used an age-of-allele test [70] to detect whether *G2/Jockey-3* frequencies deviate from our expectations. We estimated age as the number of unique substitutions for each TE copy based on the reference genome and we estimated frequency from the 28 high-coverage DGRP samples, controlling for demographic history [70].

For this test, we need to compare insertions spanning a range of ages to identify individual insertion frequencies that deviate from expectations [70]. We thus examine *G2/Jockey-3* and other TE families that have centromere insertions. We find that *G2/Jockey-3* (Figure 6B, File S9), like the majority of TEs, segregates either at the expected frequencies or lower than expected, given their age (Figure S6, File S9). This suggests that many *G2/Jockey-3* copies, both within and outside the centromeres, are either neutral or weakly deleterious. Therefore *G2/Jockey-3* shares similar selective pressures as other presumably weakly deleterious TE families in the genome.

Interestingly, we observed an enrichment of copies with zero unique substitutions for the heterochromatin and CenY insertions (Figure S5). However, we suspect this pattern arises from duplications rather than independent *G2/Jockey-3* insertion events (see below). Genomic duplication events are common in heterochromatin in the Y chromosome [73, 76, 90].

While we do not see evidence for strong recent positive selection on any individual *G2/Jockey-3* insertions in the centromeres, there may be selection on centromere haplotypes that differ in their composition or structure.

### 7. Centromere island haplotype structure not likely driven by G2/Jockey-3 insertions

To detect signatures of natural selection on centromere haplotypes and any associations with *G2/Jockey-3* copies, we examined patterns of nucleotide variation. Centromeres and pericentromeric regions in *D. melanogaster* generally do not recombine by crossing over [91, 92] therefore the effects of linked selection in non-recombining regions should lead to low levels of nucleotide diversity. We thus expect to see reduced genetic variation in the centromeres and high levels of linkage disequilibrium. We quantified genetic variation in the centromere islands for DGRP samples with high centromere coverage (> 20X, see Methods). As expected, nucleotide diversity is greatly reduced in every centromere compared to the major chromosome arms (MWU *P* < 2.0E-07; Figure S7, Table S5) except for chromosome 4, which does not recombine, and its centromere and chromosome arms are similar (MWU *p* = 0.56; Figure S8, Table S5). We find that Tajima’s *D*, a summary of the site frequency spectrum, is also typically lower in centromere islands (Figure S8), indicating a skew toward rare alleles in centromeres. The centromere island haplotype structures (Figures S9-S11) indicate that the majority of centromeric SNPs are randomly distributed with respect to the *G2/Jockey-3* sequences (FET *P* > 0.46). Linked selection anywhere in or near centromeres can reduce genetic diversity, skew frequency spectra, and create haplotype structure. While centromere island haplotype structure is consistent with a history of linked selection, we do not detect evidence that *G2/Jockey-3* insertions themselves are necessarily the targets of selection.

### 8. G2/Jockey-3 copy number amplification on the Y chromosome is driven by duplication

The Y chromosome centromere is unique in that it has the largest centromere island containing nearly half of all *G2/Jockey-3* fragments, but no identifiable satellite sequences [53]. We cannot use the genomic re-sequencing data to study polymorphic insertions on the Y chromosome, as the data are either female-only or mixed sex and thus lower coverage. However, our high-quality reference assembly provides an opportunity to infer the organization and evolutionary history of Y-linked *G2/Jockey-3*. We inferred a phylogenetic tree of all Y-linked copies of *G2/Jockey-3.* Based on this phylogeny, we classified centromeric Y-linked *G2/Jockey-3* copies into four broad classes (Figure S12). Class 1 *G2/Jockey-3* correspond to young and mostly long insertions, class 2 to short (< 160 bp) 3’ end fragments, class 3 to old internal fragments missing the 5’ and 3’ terminal ends, and class 4 to generally long and old 3’ end fragments divided into two subclasses (4 and 4A) based on position (Figure 9, Figure S12). We mapped these classifications onto our Y chromosome assembly and estimated pairwise divergence between all *G2/Jockey-3* copies in the Y centromere (Figure 7) and elsewhere on the Y chromosome (Figure S13). Each class seems to undergo independent duplication events, however at a larger scale we do not identify higher order repeat domains (Figure 7). This structure differs from other known centromeres (*e.g.* primates), which are organized into higher order repeats, the youngest repeats are found in the center and the older, more divergent repeats, are at the outer edges of the array [51, 52]. By contrast, in the *D. melanogaster* Y centromere we observe that regions enriched with young *G2/Jockey-3* copies punctuate regions rich with older copies (Figure 9), suggesting that recent, local rearrangements affect centromere organization.

**Figure 7:**
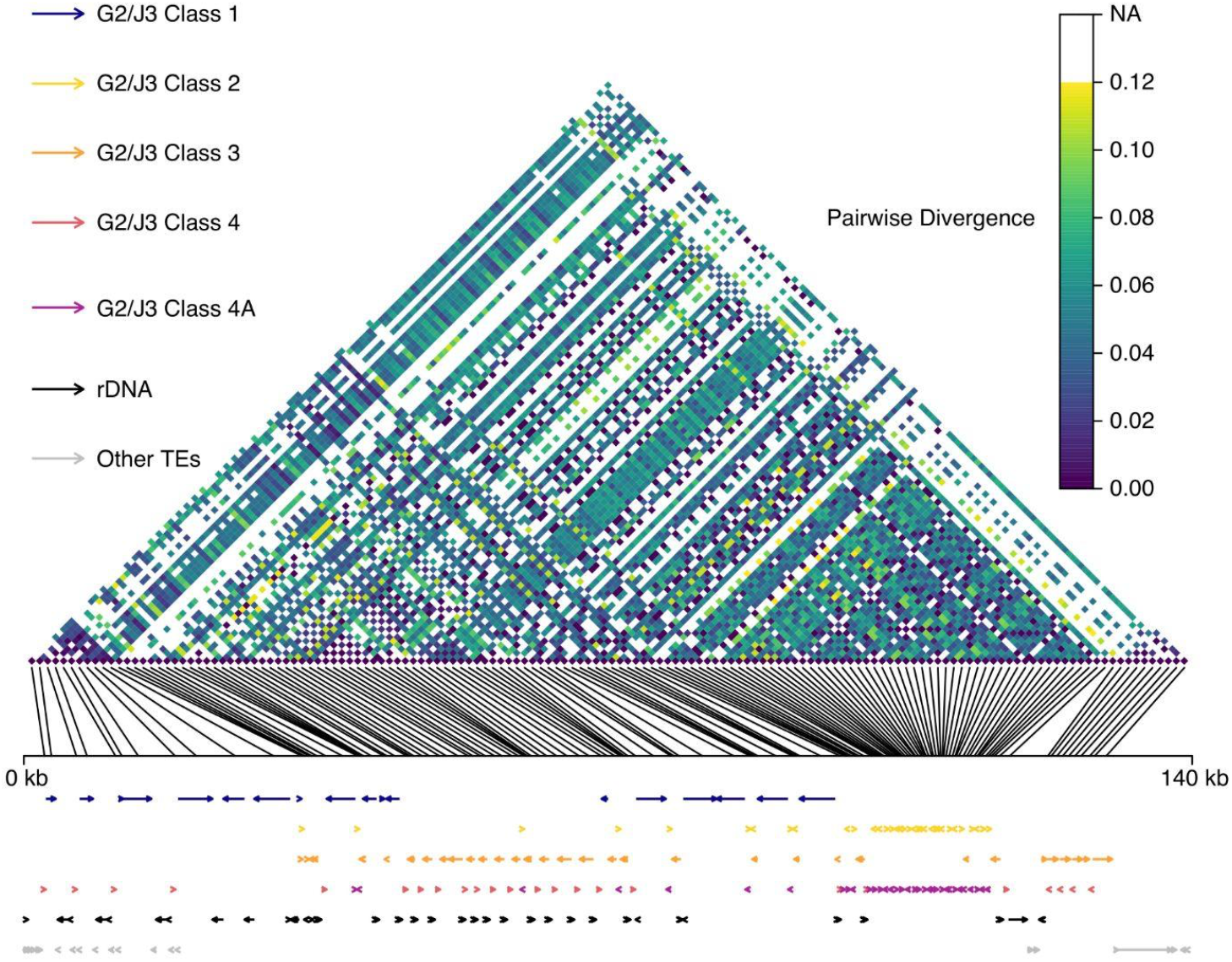
*G2/Jockey-3* evolution in the Y chromosome centromere is shaped by expansion via duplication, gene conversion and rearrangements. The colored pairwise divergences between individual *G2/Jockey-3* copies on the Y centromere are shown on top with their relative positions on the line graph below. White spaces represent *G2/Jockey-3* copies which do not overlap, so pairwise divergence cannot be calculated. Below the graph is a visual representation of the broad categories of repeats on the Y chromosome centromere, labeled by color with length and direction illustrated with arrowed lines. The repeats appear arranged into roughly four regions with overlapping borders based on the types of repeats and pairwise divergence. The classes of *G2/Jockey-3* are as follows: Class 1 younger insertions, Class 2 short (< 160 bp) 3’ end fragments, Class 3 older internal fragments missing the 5’ and 3’ terminal ends, Class 4 generally longer and older 3’ end fragments. Class 4 contains a subclade (4A) with highly similar sequences almost exclusively from the single region of CenY.

We estimated gene conversion rates of *G2/Jockey-3* on the Y centromere with respect to their classes (Supplemental Text). Our estimates for gene conversion are on the order of 1E-6 per site per generation (Table S6), ~10E3 times higher than the other chromosomes in *Drosophila* [91–93] and an order of magnitude less than previously estimated gene conversion rates between duplicated gene families on the Y chromosome [73]. Moreover, we do not see complete sequence homogenization within classes. We observe similar patterns of duplication and expansion outside the centromere on the Y chromosome (Figure S13), as 32 of 47 non-centromere Y-linked copies appear to be duplicated; they are of similar fragment size, orientation, and have few no zero unique substitutions (File S6). Therefore, we conclude that *G2/Jockey-3* evolution on the Y chromosome is driven by duplication of historical insertions with some gene conversion among copies. Population genomic data from the Y chromosome will help distinguish between types of recombination affecting Y centromere structure.

Outside of the Y chromosome, putative duplications involving *G2/Jockey-3* are less common (Figure S13). Many copies of *G2/Jockey-3* within the centromeres are fragmented and therefore share no homologous sequences for pairwise calculations (Figure S13). Pairwise divergence between centromeric *G2/Jockey-3* fragments is elevated in comparison to fragments in both the euchromatin and non-centromeric heterochromatin (Figure S14, MWU, *P* < 0.01), and lowest in non-centromeric heterochromatin, especially on the Y chromosome. Therefore, the majority of *G2/Jockey-3* within the non-Y centromeres appears to be the result of recurrent past insertions.

## CONCLUSIONS

*G2/Jockey-3* is a unique TE in that it is the only sequence shared among centromeres of *D. melanogaster* [53]. Here we show that *G2/Jockey-3* is a recently active non-LTR retroelement that may have had an insertion preference for centromeres, given their relatively small size compared to the rest of the genome (Figures 3). While the centromere enrichment and skew toward high frequency centromere copies are expected for insertions favored by natural selection, we do not find evidence for positive selection on individual *G2/Jockey-3* insertions. Instead, most *G2/Jockey-3* insertions both inside and outside of the centromeres are at the expected frequency or significantly rarer than expected given their age, suggesting they may be neutral or slightly deleterious and accumulating due to relaxed constraints against centromeric TE insertions (Figure 6B). The majority of *G2/Jockey-3* copies within the centromeres appear to be the result of recurrent past insertions, except for on the Y chromosome. Some Y-linked copies are likely a result of duplications rather than *de novo* insertions, similar to patterns reported in other species (*e.g.* [28]).

We propose that most *G2/Jockey-3* insertions in the centromeres resulted from a burst of activity, which is consistent with the evolutionary history of other non-LTR retroelements [88]. When a TE first invades, it can proliferate rapidly until it comes under the surveillance of germline defense mechanisms like the piRNA pathway, for example by inserting into a piRNA cluster [94–96]. These dynamics are consistent with the invasion and control [94, 97] of the *P* element in *D. melanogaster* [98] and its close relative, *D. simulans* [99]. Once targeted by the piRNA pathway for transcriptional and posttranscriptional silencing, the TE may eventually become dormant or go extinct. Here we show that *G2/Jockey-3* had a burst of activity and is now inside piRNA clusters and actively targeted by the piRNA pathway (Table S4).

The *G2/Jockey-3* insertion polymorphism patterns and piRNA observations are most consistent with some underlying selfish behavior typical among TE families, except that this element can target centromeres. Although we are unsure of the precise targeting mechanism, we suspect the target is the centromere chromatin rather than any particular DNA sequence. Such centromere targeting was reported for LTR retroelements in other species such as maize [100, 101] and *Arabidopsis* [102, 103]. For example, when the *Arabidopsis lyrata Tal-1 Copia-type* retroelement [103] is introduced into *A. thaliana*, the element targets the centromeres despite divergence in centromeric DNA sequences between these species [102].

TEs may accumulate in heterochromatic regions where they experience relaxed selection compared to euchromatic insertions, which have potential to disrupt functional elements, risk ectopic exchange and resulting genomic rearrangements [85, 86, 104], and spread silent chromatin or DNA methylation into nearby functional elements [*e.g*., 105–108]. However, centromeres are not typical heterochromatin [109]. Here we propose that centromeres offer a unique environment for TEs (Figure 8A). First, insertions are unlikely to be strongly disfavored, as the function of centromeres is unlikely to be disrupted by a single insertion event. Second, centromeric insertions may escape host silencing mechanisms. Small RNAs target TEs to limit potentially harmful TE activity by inducing H3K9 methylation and thus transcriptional silencing (reviewed in [110–112]). However, CENP-A structure and posttranslational modifications differ from H3 [109], and the host silencing machinery is not known to target CENP-A. Therefore, centromeres may offer a ‘safe haven’ for some TEs not just because they can avoid removal by purifying selection, but also because TEs may temporarily escape targeting by small RNAs (*e.g.* piRNAs, siRNAs) and alternative host defenses (*e.g.* DNA methylation in pericentromeres [113], hypomethylated DNA in humans [52, 114]).

**Figure 8.**
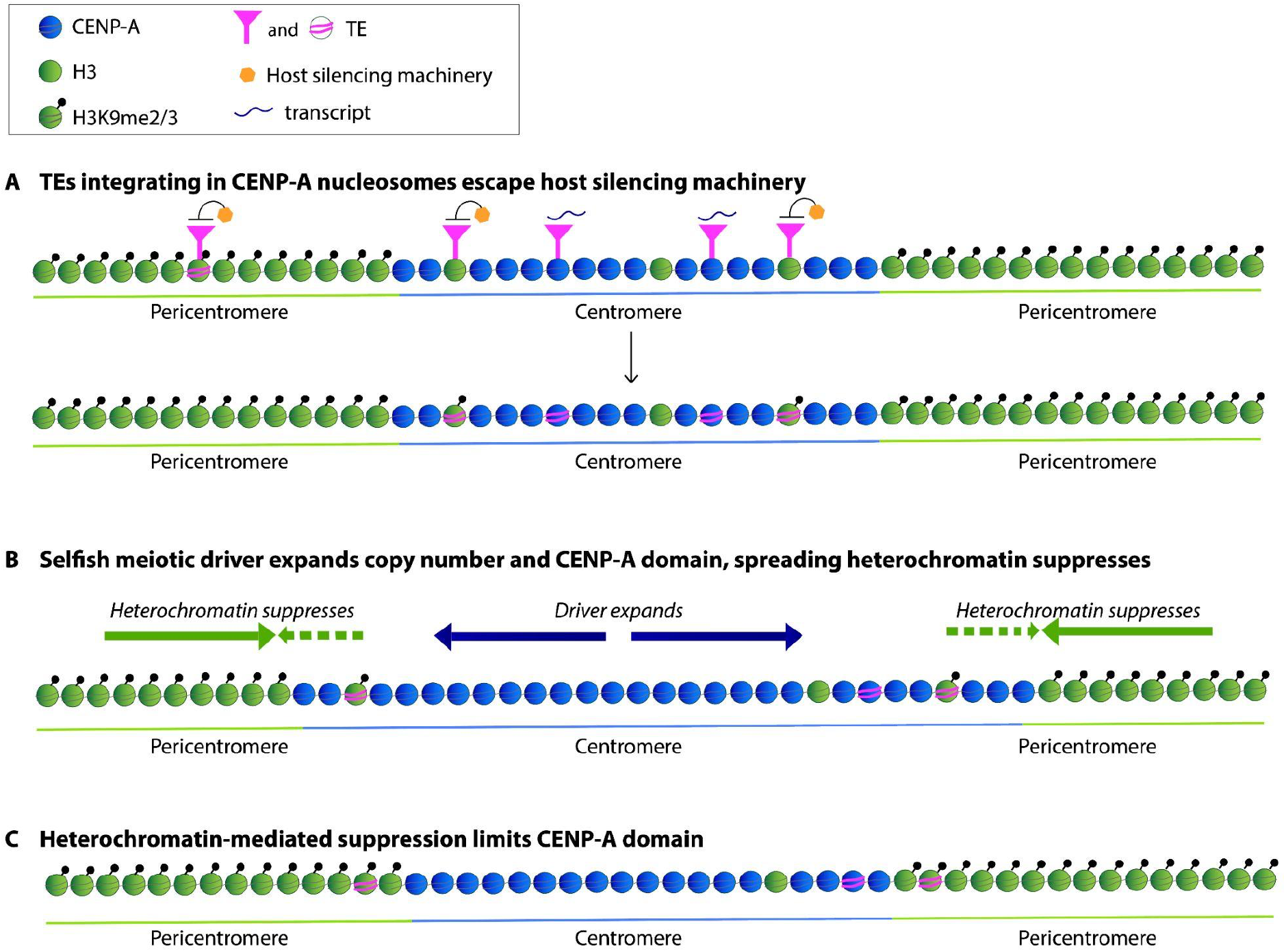
Model for the accumulation of centromere-associated retroelements. Centromeres have unique chromatin with interspersed CENP-A and H3 domains, including some H3K9me2/3 [12–15]. The diagrams represent nucleosomes, either CENP-A (blue) or H3 (green), with H3K9me2/3 modifications indicated with a black dot. The nucleosome distributions are not drawn to scale. A. Active TEs like *G2/Jockey-3* jump in centromeric chromatin (magenta triangles and lines in nucleosomes) and could contribute to transcription (black wavy line) or be silenced by host machinery (orange hexagon). Centromeres may offer a safe haven for TE insertion because insertions there are unlikely to be strongly disfavored and because insertions in CENP-A nucleosomes may escape host silencing machinery (*e.g.* piRNA pathway). Centromeric TE insertions might be tolerated or even favored by the host if they contribute to centromere transcription or a low level of H3K9me2/3 inside centromeres. B. Selfish centromeric DNA sequences can expand their copy number and bias their transmission to the next generation through female meiosis [131]. C. One way to suppress cheating centromeres is by spreading heterochromatin to limit the CENP-A domain [136]. TE insertions in H3 nucleosomes within the centromeres may be targeted for silencing by host silencing machinery (green circle with magenta lines and black circle). These H3K9me2/3 nucleosomes within centromeres may help create a more efficient response to driving centromeres by merging spreading pericentric heterochromatin with internal H3K9me2/3 domains.

While most TEs are considered to be parasitic elements that are neutral or mildly deleterious [115, 116], some TE insertions, including those in non-recombining and/or gene poor heterochromatic environments, might contribute to host function [117]. For example, TEs can adopt commensal and/or mutualistic strategies to propagate without direct confrontation by cooperating with their host (reviewed in [118]). One extreme outcome is TE domestication, where the elements no longer pose a threat to the host and instead now perform some important genome function. One often cited example is at *Drosophila* telomeres, where three non-LTR retroelements in the *Jockey* clade (*HeT-A, TAHRE*, and *TART* collectively referred to as HTT) have specialized roles [119, 120]. The transposition activity of HTT is limited to telomeres, where they directly contribute to telomere length and integrity [119–122]. However, detailed study of the evolutionary genetic history of the retroelements across species suggests instead that there is still an underlying conflict between the HTT and the genome, as there is rapid turnover of HTT between species [123]. Alternatively, given the specialization of HTT, the dynamic and unstable telomeric environment may necessitate rapid HTT evolution [124].

While centromeric TEs do not currently show evidence for domestication in *D. melanogaster*, they may be tolerated, or even beneficial for three reasons. First, TEs may contribute to centromere transcription (reviewed in [35]), as some transcription at centromeres is important for CENP-A assembly and function [14, 46, 125–129] and centromere function [14, 45, 130]. Second, TEs might contribute to the unique chromatin at centromeres. H3 nucleosomes are typically interspersed with CENP-A within centromere domains [12, 14] [15], including a limited amount of H3K9me2/3 (*e.g.*, [13]). We propose that insertion of active TEs like *G2/Jockey-3* may contribute to this unique CENP-A/H3 chromatin at the centromeres (Figure 8A). While landing in CENP-A nucleosomes may allow the TE to temporarily escape silencing, if an active TE lands in an H3 nucleosome region of the centromere, it may be targeted for silencing through H3K9 methylation (*e.g.* piRNA- or siRNA-mediated silencing) and thus contribute to H3K9me2/3 domains within the centromere. Third, TEs may help mitigate other conflicts that happen at centromeres. Some centromeric sequences have the capacity to be selfish and bias their transmission through female meiosis in a process known as meiotic drive [131]. Driving centromeres generally have expansions of satellite DNA sequences that can recruit more kinetochore proteins and cause biased segregation toward the egg pole [132, 133] (Figure 8B). Meiotic drive at the centromeres can lead to rapid evolution of both centromere DNA and proteins that work to restore fair segregation [134, 135]. There are parallel pathways to suppress meiotic drive involving rebalancing kinetochore proteins and spreading heterochromatin to limit the CENP-A domain [136]. While centromere drive and suppression models predict that the flanking heterochromatin domains are the main participants in the conflict [136], the H3K9me2/3 domains within the centromeres might also contribute. Active centromeric TEs that are targeted by small RNA pathways, like *G2/Jockey-3*, could lead to spreading of H3K9me2/3 heterochromatin from within the centromere (Figure 8C). In this way, the centromeric TEs and ‘internal’ centromeric H3K9 domains may also be a substrate for suppressors of centromere drive to stop the unchecked spread of CENP-A caused by meiotic drivers.

Even if TEs cooperate with centromeres, this may be an uneasy alliance, as there could still be an underlying conflict. For example, the invasion of H3K9me2/3 heterochromatin in centromeres is potentially dangerous, as it may evict too much CENP-A and silence a centromere [5]. Therefore, the centromeric chromatin, and the relative abundance of CENP-A and H3 nucleosomes, may mediate the balance between conflict and cooperation between active TEs and centromeres.

In many species across the tree of life, centromeres are associated with TEs [18, 23–34, 53]. The connection between TEs and centromeres is supported not just by TE sequences embedded in and around present-day centromeres, but also in the origins of some proteins that function in centromere biology (*e.g.* similarity between centromere protein CENP-B and Pogo family transposons [137]). This relationship may represent a delicate balance between conflict and cooperation between the centromeres and the repetitive selfish genetic elements that occupy these genome regions. We expect this balance to vary depending on genomic context, level of TE activity, and presence and frequency of centromere meiotic drive. If TEs contribute positively to centromere function, many types of elements may be sufficient to provide the right ‘environment’. In this case, we would not necessarily expect to see a signature of selection on any one insertion or type of insertion. While *G2/Jockey-3* is currently the principal element occupying the centromere niche in *D. melanogaster*, it may be replaced over time by another type of repeat. In our model, any repeat that can be transcribed and can recruit H3K9me2/3 (*e.g.* is targeted by small RNAs) can contribute to centromere function. Detecting the mode, tempo, and target of selection in non-recombining and dynamic environments like the centromeres is difficult. Exploring the evolution of centromere composition in closely related species will provide additional insights necessary to test these hypotheses.

## MATERIALS AND METHODS

### G2/Jockey-3 re-annotation and expression analysis

Chang et. al. [53] generated the *G2/Jockey-3* consensus sequence after a phylogenetic analysis confirmed previously annotated *G2* and *Jockey-3* were the same element [53]. We annotated a heterochromatin-enriched long-read genome assembly of *D. melanogaster* [73] with Repeatmasker v4.1.0 using the previous *G2/Jockey-3* consensus sequence along with a *Drosophila* repeat library [138]. We extracted *G2/Jockey-3* and the surrounding sequences using BEDTools v2.29.2 [139], aligned the sequences to the previous consensus sequence, manually inspected the alignment, and generated the new consensus sequence with Geneious v8.1.6 [140] (File S1). We identified Open Reading Frames (ORFs) with NCBI ORFfinder (https://www.ncbi.nlm.nih.gov/orffinder/) and screened the ORF candidates by identifying conserved domains common in non-LTR retrotransposons with the NCBI Conserved Domains search tool (https://www.ncbi.nlm.nih.gov/Structure/cdd/wrpsb.cgi). We defined the 5’ end by identifying transcription along the length of *G2/Jockey-3* by mapping publicly-available single-end Precision nuclear Run-On Sequencing (PROseq) downloaded from NCBI SRA under BioProject PRJNA451204 and accessions SRR7050623 - SRR7050630 [62]. We removed adapters and reads < 16bp using cutadapt v2.1. Reads were reverse complemented using the fastX toolkit v0.0.13 (fastx_reverse_complement). We mapped reads to the heterochromatin-enriched genome assembly of *D. melanogaster* using bowtie2 v2.3.5.1 (--no-discordant -- dovetail --very-sensitive options; [141]). We extracted reads > 16 bp in length mapping to *G2/Jockey-3* annotations in the genome with samtools v1.7 [142] and remapped the reads to the *G2/Jockey-3* consensus with bowtie2 v2.3.5.1 [141]. We separated reads based on their strand using samtools (v1.7). We calculated read depth at each position along the *G2/Jockey-3* consensus and normalized coverage by the total number of genome mapped reads (RPM) using bedtools genomecov [139]. We used alignment depth of all *G2/Jockey-3* genomic fragments to identify where transcription occurs along the length of *G2/Jockey-3*. We submitted the sequence in 5’ end motifs under transcription peaks for a GoMo analysis [143] to identify motifs as possible promoters.

To detect piRNAs corresponding to *G2/Jockey-3*, we trimmed the small RNA-seq reads for adaptors and excluded the reads that align to rRNA or tRNA. We aligned the remaining reads to the heterochromatin-enriched genome assembly [53, 73] using Bowtie [144], and then used a customized python script as described in Wei et al. [83] to count reads that mapped to each TE feature (htseq_bam_count_proportional.py; https://github.com/LarracuenteLab/Dmelanogaster_satDNA_regulation; last accessed 11/2022, https://doi.org/10.5061/dryad.hdr7sqvj3). We reported the RPM (Reads Per Million) values in a summary table. We calculated the 10nt overlap Z-score of piRNAs using piPipes [145].

### Short-read analysis

We detected TE insertions from publicly available FASTQ files from 135 *Drosophila melanogaster* Genetic Reference Panel (DGRP) lines from BioProject PRJNA36679 [64, 65] downloaded from the European Nucleotide Archive last accessed September 20, 2019, 13 of the founder lines of the Drosophila Synthetic Population Resource (DSPR) BioProject PRJNA156883[146], and 85 samples from the Global Diversity Lines BioProject PRJNA268111 (Table S3)[147]. We detected TE insertions in each sample with the McClintock pipeline [69] (last downloaded and updated, August 20, 2020) combining several split-read and discordant read mapping programs including the following: ngs_te_mapper, popoolationTE, popoolationTE2, Retroseq, TE-locate, and TEMP (scripts and files for running McClintock at https://github.com/LarracuenteLab/Dmel_Jockey-3_Evolution). We concatenated the outputs to calculate the number of TEs detected by each program and sample and analyzed the outputs with Python and R scripts available at https://github.com/LarracuenteLab/Dmel_Jockey-3_Evolution. We repeated our analyses of TEs with and without insertions in known piRNA clusters and removed fragments of Gypsy6_I_Dmoj and Gypsy1_I_Dmoj which resemble satellite sequences in *D. melanogaster.* Depth of each short-read sample was calculated from the sorted BAM outputs from the McClintock pipeline using Mosdepth [148] (Table S3). We retained samples which had a centromere coverage greater than 20X for our analyses (see Supplemental Text for more information).

We used the same short-read FASTQ files from the DGRP to quantify genetic diversity between the centromere islands and the rest of the genome using a best practices GATK pipeline v4.1.9.0 to detect variant SNPs. We mapped reads from high coverage samples to our reference genome with BWA v1.7.15 and samtools 1.7 to generate sorted BAM files and used GATK MarkDuplicates, BaseRecalibrator, ApplyBQSR, AnalyzeCovariates, and HaplotypeCaller to call and filter SNPs for each sample into a VCF file. The VCFs for all samples were concatenated with GenotypeGVCFs and relicabrated with VariantRecalibrator and ApplyVQSR. We quality filtered SNPs in the resulting GVCF file with the following criteria; QD < 2.0, FS > 60.0, MQ < 40.0, MQRankSum < −12.5, and ReadPosRankSum < −8.0. We calculated *π* and Tajima’s *D* with the SimplifyVCF_Basic.v2.pl and Windows_Basic.v2.pl Perl scripts on the filtered SNPs (available at https://github.com/bnavarrodominguez/SD-Population-genomics, [149]). We further analyzed variants within the centromeres with VCFtools [150] and the vcfR R package [151]. Additional details of our analysis with short-read data and the results can be found in the Supplemental Text section.

### Long-read analysis

We assembled genomes for several *D. melanogaster* lines following the heterochromatin-enriched assembly protocol from Chang & Larracuente [73]. We downloaded raw PacBio reads for five of the Global Diversity Lines (GDL) from the NBCI BioProject PRJNA451471 [68], eight Drosophila Synthetic Population Resources (DSPR) founder lines and Oregon-R with raw Pacbio reads from BioProject PRJNA418342 [67], and five DGRP lines from Bioproject PRJNA559813 [66]. We assembled draft genomes using all reads with Canu v1.7.1 [152] with the relaxed corrected error rate and the parameters “genomeSize=160m useGrid=false stopOnReadQuality=false corMinCoverage=0 corOutCoverage=100 correctedErrorRate=0.100”. We then mapped PacBio reads to the reference genome with minimap2 (downloaded August 28, 2018) with the “-x map-bp” parameter setting [153]. We then used unmapped reads and reads mapping to heterochromatic regions to generate an additional “heterochromatin-enriched” assembly. We merged the two assemblies into a single assembly with quickmerge [154] and polished the assemblies with Racon v1.3.1 [155] by mapping the PacBio reads three times to the assembly using minimap2 [153]. We also polished the GDL assemblies twice with Illumina reads from BioProject PRJNA268111 [147], the DSPR assemblies with reads from https://wfitch.bio.uci.edu/~dspr/index.html (last accessed September 6, 2021 [156]), BioProject PRJNA252961 for Oregon-R, and the DGRP assemblies with reads from BioProject PRJNA36679 [64, 65]. We annotated all assemblies with Repeatmasker v4.1.0 and our *Drosophila* TE library with the updated *G2/Jockey-3* consensus sequence [138] (File S10). We identified centromere island sequences in assemblies using a combination of methods to find sequences corresponding to the centromere islands identified in our reference genome [53]. These include aligning assemblies to the centromere islands with MUMMER v.3.23 nucmer [157], identification of similar sequences with BLAST 2.10.0+ [158], and identifying contigs enriched in *G2/Jockey-3* or satellite DNA repeats enriched around the centromere islands [53]. We also used MEME [159] to detect motifs enriched in the flanking regions of young insertions in our assemblies. Additional details of our analysis with long-read data, the results, and the MEME motif analysis are in the Supplemental Text.

### Insertion polymorphism validation

We validated TE insertion predictions in three ways: 1) we used short reads to confirm TE predictions in long read assemblies (see Supplemental Text); 2) we used long read assemblies to confirm TE predictions in short read data (see Supplemental Text); and 3) we selected a subset to validate with PCR (File S3). We used the results from these analyses to establish criteria for including a predicted insertion in our analyses. For PCR, we selected a subset of 59 *G2/Jockey-3* copies to confirm in four GDL lines; B59, I23, T29A, and ZH26. We included copies from each chromosome (X, 2, 3, 4), each chromatin region (euchromatin, heterochromatin, and centromere), and both reference and non-reference copies. We were able to detect 33 copies including 13 copies in the centromere and seven *de novo* insertions (File S3).

We used permutation tests to test for significant differences in density of recent *G2/Jockey-3* insertions between genome regions. For these tests, the genome was divided into 90-kb windows (average size of a centromere, excluding the Y chromosome and 2^nd^ chromosome) and categorized as centromere, heterochromatin, or euchromatin. We estimated *G2/Jockey-3* counts in each window based on final insertion polymorphisms detected by short-read and long-read datasets separately. We excluded the Y chromosome from these analyses. To test for enrichment of *G2/Jockey-3* counts within categories, we permuted the datasets 10,000 times using regioneR 1.18.1 ([160] ntimes=10000, randomize.function=resampleRegions, evaluate.function=numOverlaps). Results were similar for permutation tests with the whole genome, excluding heterochromatin windows, and excluding euchromatin windows.

### G2/Jockey-3 sequence and phylogenetic analysis in D. melanogaster

We extracted all *G2/Jockey-3* sequences from the annotated assemblies using BEDTools v2.29.2 [139] and classified the insertions as reference or non-reference (*de novo*). We classified insertions as homologous to those in our reference if they were both fragments of similar size (+/- 10 bp) and within +100 bp of the same location in the reference genome. We identified corresponding locations by aligning the assemblies to the reference genome with nucmer within MUMMER v.3.23 [157]. Insertions were classified as *de novo* if they differed in fragment size length and position of reference insertions, found outside the reference fragment position range, and located within breaks in the nucmer alignment to the reference genome.

We performed phylogenetic analyses for candidate *G2/Jockey-3* sequences identified in our genome assemblies. We used *G2/Jockey-3* sequences from the reference and the non-reference insertions discovered in the DSPR and GDL assemblies. If a non-reference insertion was found in multiple assemblies, we randomly selected an assembly from which to extract that insertion to include in our analysis. Prior to phylogenetic analysis, we filtered sequences to exclude fragments shorter than 400 bp or ~10% of the length of a full *G2/Jockey-3* element. We aligned the sequences with MAFFT [161] with the *D. simulans G2/Jockey-3* consensus as the outgroup and inferred a maximum likelihood phylogenetic tree with RAxML v8.2.11 [162]. To study the evolution of *G2/Jockey-3* on the Y chromosome, we inferred a maximum likelihood phylogeny of all fragments exclusive to the Y chromosome regardless of size. We determined the relative age of individual insertions using divergence from the consensus from the Repeatmasker output and the number of substitutions unique to a fragment divided by fragment length. “Young” insertions were either < 1% diverged from the consensus sequence or *de novo* insertions exclusive to a single genome assembly and < 2% diverged from the consensus sequence. We calculated pairwise divergence between fragments with MEGA X [163] using the Kimura 2-parameter model assuming uniform rates among all sites. We estimated gene conversion rates between elements using the same method outlined from Chang & Larracuente [73] based on Ohta’s a equation [164] and Rozen’s equation [165]. We used the following equation for our calculation of gene conversion:

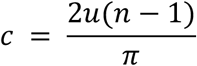

We determined the rate at which a number of sequences (*n*) homogenize each other per generation (*c*) based on the point mutation rate (*u*) and divergence among the sequences (+). We grouped reference *G2/Jockey-3* sequences located within the same contig except for insertions on the Y centromere. Fragments from the Y centromere were grouped based on the phylogenetically-defined classes. We aligned the sequences with MAFFT [161] and calculated divergence with MEGA X [163]. We normalized *c* based on gene conversion tract length of a single event (*c_g_*) based on previously estimated 400 bp tract length of a single event in *D. melanogaster* [92, 93]:

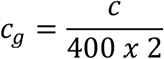

### Age-of-allele test

Natural selection, transposition, and frequency of insertion influence TE evolution within populations. We used an age-of-allele test [70] to determine departure from neutral evolution of individual insertions based on population frequency, age, and demographic history in *D. melanogaster.* The age-of-allele test determines the probability distribution of an insertion frequency within a population under neutrality conditioned on the age of the insertion. The age is inferred by the number of unique substitutions within a TE sequence accumulated since the time of the insertion. To determine the number of unique substitutions, we aligned insertions with MAFFT [161] for *G2/Jockey-3, BS, G5, Doc*, and *Doc2* detected as reference insertions in the McClintock output. Unique substitutions were quantified from each individual insertion using Biostrings [166]. We calculated the probability distribution of the frequency of each unique insertion with the age-of-allele test script from Blumenstiel et al. [70] in Mathematica v12.0.0.0. The probability distribution is also conditioned on population demography of North American *D. melanogaster* populations outlined in the original paper [70]. North American *D. melanogaster* populations experienced two bottlenecks in their migration from Africa [167, 168] that led to non-equilibrium TE evolution [70]. We compared the probability distribution to the frequency of insertions detected in the 28 DGRP samples with >25X average coverage of the centromere islands. We excluded insertions in the Y chromosome given the high rate of duplication. The expected frequency of each insertion in the population is the probability of observing an insertion as frequent or more frequent within the population multiplied by the observed number of insertions in our population sample. We calculated significant deviations from neutrality as probability of observing as many or fewer, or as many or greater, copies < 0.1.

## DATA AVAILABILITY STATEMENT

All scripts for data analysis and data for figure generation are available on Github at https://github.com/LarracuenteLab/Dmel_Jockey-3_Evolution. The genome assemblies, genome annotations, and large datasets including the McClintock outputs will be available on Dryad (DOI forthcoming).

## ACKNOWLEDGEMENTS

This work was funded by the National Science Foundation (NSF MCB 1844693) and University of Rochester funds to AML. We thank the Mellone lab, Dr. Justin Blumenstiel, and the Larracuente lab for helpful discussion. We also thank the University of Rochester Center for Integrated Resource Computing for access to computing cluster resources.

