## Supplemental text and figures for "Centromere-associated retroelement evolution in *Drosophila melanogaster* reveals an underlying conflict"

*Summary of short- and long-read data*

To investigate the evolutionary history of *G2/Jockey-3* and other TEs within and outside the centromeres, we surveyed available whole genome shotgun *D. melanogaster* genomic population resources (Table S1), excluding any datasets generated with whole genome amplification. The short-read datasets include many population samples for surveying patterns of TE insertion polymorphisms. We surveyed a sample of lines from short-read datasets of the Global Diversity Lines (GDL) [1] and the founder lines of the Drosophila Synthetic Population Resource (DSPR) [2] as well as a deeply sampled North American population from the Drosophila Genetic Reference Panel (DGRP) [3, 4]. Highly repetitive regions of the genome are typically underrepresented in short-read data [5, 6]. However, the centromere islands, while embedded within satellite DNA, contain unique DNA sequences. We determined that the coverage of the centromere islands is consistently lower than the euchromatic regions of the genome. (Mann Whitney U $P = 1.46E\text{-}10$, Table S1, Figure ST1 and ST2). Mean genomic coverage in the GDL samples was consistently lower (~12X) than both the DGRP and DSPR (~33X and ~60X respectively, Figure ST1 and ST2, Mann Whitney U, $P < 1E\text{-}7$); the median coverage of the centromere islands were ~6X, ~17X, and ~26X respectively. The centromere coverage in the GDL short-read dataset is too low to accurately detect TE insertions. Samples within the DGRP were also variable in genomic coverage. Therefore, we only included samples which had a median centromere coverage of 20X or greater to stay within range of the DSPR lines.

We detected *G2/Jockey-3* and other TE insertions using the McClintock meta-pipeline combining several TE detection programs [7] and filtered out calls which were either redundant or mislabeled. After filtering, there were 949,097 total TE calls detected by the programs within the pipeline corresponding to 47,574 unique TE insertions (File S4, File S5). Of the unique TE insertions, 50% (23,636 / 47,574) were new insertions not found within our reference genome assembly. Most TEs are presumed to be deleterious and therefore expected to be segregating at low frequencies within the population [8]. We find 38% (18,248 / 47,574) of all TEs within the DGRP are detected within just one sample.

In addition, we generated *D. melanogaster* genome assemblies with PacBio long reads from a global sample of flies from almost every continent (Table 1). These include five samples from the GDL [9], 13 of the (DSPR) founder lines, Oregon-R [10], and five DGRP lines [11]. Although the samples do not provide a deep insight into the evolution of TEs in centromere islands from a single population, they provide the sequence data to directly calculate the age and abundance of TE among a global sample of *D. melanogaster*. The long-read assemblies provide more comprehensive coverage of TE dynamics in heterochromatic regions including the regions corresponding to the centromere islands identified by CENP-A ChIP-seq in the ISO1 reference strain [12]. We were able to identify almost all of the sequences corresponding to the centromere islands on chromosomes X, 3, and 4 in the assemblies (Table S2). Cen2 was absent from the DGRP and DSPR datasets and one DGRP assembly. It is difficult to sequence Cen2 because with a length of 20 kb with flanking satellite sequence it is small even compared to the other centromeres. The Y chromosome was also notoriously underrepresented from the long-read datasets except for tiny fragments. This is most likely due to reduced coverage of the haploid Y chromosome from sequencing a mix of males and females for the GDL population samples [9]; only females were sequenced for the DSPR founder lines [10] and the DGRP lines [11].

*Reliability of TE detection using short reads.*

TE detection in short-read data is notoriously difficult because mapping reads to individual TEs and detecting *de novo* elements is a challenge [13]. We compared TE detection in two ways. First, we determined what proportion of the *G2/Jockey-3* copies in the genome assemblies were detected in short reads by McClintock. This was done to determine how reliable McClintock was at detecting *G2/Jockey-3* in different regions of the genome, especially the centromeres. Second, we analyzed all of the *G2/Jockey-3* copies detected by McClintock to see if these were found in the assembly. This analysis was done to determine the proportion of true positives to false positives detected by McClintock. A TE detected by McClintock and found in an assembly was counted as a true positive. We considered false positives to be TEs called by the McClintock pipeline but were confirmed missing from the assembly. We used nucmer within MUMMER v.3.23 [14] to align the genome assemblies to our reference genome and identify the sites of *G2/Jockey-3* according to the reference (File S3). Then we checked for overlap between the copies detected in our assembly annotations completed by RepeatMasker v4.1.0 and the McClintock pipeline. Some regions with repetitive sequences were difficult to accurately identify the corresponding region in the reference genome and were left "unknown."

We first investigated concordance between TEs detected by McClintock and the genome assemblies for the founding DSPR lines (Table S1, File S2, File S3). The average genome coverage of the short-reads for DSPR lines was greater than the other two sources (Figure ST1, Figure ST2). In the assemblies, we detected approximately twice as many insertions regardless of TE family in the long-read assemblies in comparison to the McClintock pipeline (Figure ST3). This is most likely due to the number of elements detected in heterochromatic regions in the long-read read assemblies where mapping short reads proves difficult. For reference insertions of *G2/Jockey-3*, McClintock was unable to detect approximately two-thirds of the insertions from the assemblies (123 / 357); approximately half of the reference copies were undetected by McClintock in both the euchromatin and centromere islands (152 / 280 and 287 / 535 respectively). Most of the reference copies missed were either small (<100 bp) or disconnected pieces of older, broken copies. Meanwhile, McClintock detected the majority of *de novo* copies in the DSPR assemblies (108 / 172). A majority of the missed *de novo* copies were in the heterochromatin (33 / 72). McClintock was better at detecting most of the *de novo* copies in the euchromatin (93 / 120) and the centromere (7 / 10). We next examined all McClintock calls and determined how many calls corresponded to

TEs detected in the long read assemblies. This was used to determine the ability of the pipeline to detect true positives and calculate a false positive rate. The pipeline was reliable at detecting reference TEs in the euchromatin and centromeres. 96% (367 / 384) of McClintock calls, including 95% (151 / 159) of copies in the centromere, were identified in the assemblies. We confirmed an additional 11 McClintock calls in the heterochromatin regions in the assemblies but could not confirm 49 calls; the regions were either missing from the assembly or difficult to place within highly repetitive regions. For *de novo* insertions, McClintock was successful at detecting most *G2/Jockey-3* copies in euchromatin (57 / 71) and the centromere (4 / 7) while performing worse in detecting insertions within non-centromere heterochromatin (6 / 21). However, McClintock on stringent settings detected an additional 273 non-reference copies not found in the assemblies. Out of the copies missing, 55 were in highly repetitive regions which did not align well to the reference genome for confirmation. We inspected the regions which aligned to the reference genome and confirmed the remaining 218 non-reference calls were missing from the assemblies. Out of all of the non-reference that were missing, 212 were only detected by a single McClintock method and 189 were detected in a single sample. Only six of the *de novo G2/Jockey-3* copies detected by one method and or 36 copies detected in multiple samples were found in the DSPR assemblies. The false positive rates plummet when multiple methods within the McClintock pipeline detect the same *de novo* copies or detect a copy in more than one sample. Only 9% (6 / 69) of copies detected by multiple methods were true false positives. Meanwhile, less than half (29 / 66) of insertions detected in multiple samples were false positives.

We assembled genomes for five DGRP lines from publicly available PacBio data [11] which were then annotated with our TE libraries. These were compared to the DGRP short-read data analyzed by the McClintock pipeline. In comparison to the DSPR lines, there were fewer reads mapped to available DGRP assemblies, ranging from 5X to 18X coverage in the centromeres (Table S3). Approximately a third of the reference *G2/Jockey-3* copies from the assemblies were detected by McClintock (211 / 641), which was consistent with the DSPR results. By region, centromere reference copies were significantly depleted (26%, 86 / 328) in comparison to the heterochromatin and euchromatin with approximately 40% of insertions detected by McClintock (83 / 210, 42 / 104 respectively). Once again, most of the reference copies missed by McClintock were either small or disconnected pieces of older, broken copies. For *de novo* copies in the assemblies, McClintock detected roughly two-thirds (51 / 74) which was similar to the DSPR. The single *de novo* centromere copy in the assemblies was detected by McClintock. Once again, most of the *de novo* copies within the heterochromatin were missed by McClintock (4 / 14 detected) while the majority of the euchromatic copies were detected (45 / 58). Once again, we examined all of the McClintock calls as well to determine how many correspond to copies of *G2/Jockey-3* in the DGRP assemblies. Almost all McClintock calls for reference *G2/Jockey-3* copies were found in our assemblies *G2/Jockey-3* (105 / 106) including all 45 centromere copies; roughly half of the McClintock *de novo* calls were found in our assemblies (37 / 66). Only 2 out of 7 *de novo* calls in the centromeres were missing from the assemblies including 3 calls missing on CenX in strain 855. Nearly all of the *de novo* McClintock calls missing from the assemblies were detected by a single program (21 / 29). However, all centromere *de novo* insertions were detected by a single program.

Altogether, McClintock is reliable at detecting *G2/Jockey-3* with the appropriate filters in place. Detecting reference copies was not difficult for the pipeline outside of the smaller and older fragments. However, McClintock detects much more *de novo* copies than we can find in the assemblies. Most of these false positives are detected by a single program within the pipeline and this effect is exacerbated with the centromere copies, which are usually detected by a single program. Single programs detect these insertions about half the time in the DSPR with higher centromere coverage but less in the DGRP with lower coverage. Therefore, we kept our analysis stringent based on the following criteria. First, the short-read mapping coverage to the centromeres needed to exceed 20X. Second, all *de novo* copies needed to be confirmed with either more than one program within McClintock or by more than one sample within the population. Using these criteria in the McClintock data, we observed a decrease in the number of missing *G2/Jockey-3* copies from in the DGRP. As a reminder, none of the short-reads DGRP short reads corresponding to our assemblies met the 20X coverage criteria. The number of missing copies in the DSPR which met the coverage criteria decreased from 229 to 38. Therefore, we are confident in the McClintock calls remaining after filtering and acknowledge some error remains when detecting TEs from short-read data.

*Motif analysis of G2/Jockey-3 insertion sites*

We asked if there was any evidence for *G2/Jockey-3* target site preferences by using MEME [15] to detect motifs enriched in the flanking regions of young insertions. We extracted 100 bp upstream and downstream of each insertion and used MEME to identify enriched motifs. We detected an enrichment of rDNA-associated sequences including 18S and non-transcribed spacers (NTS) (File S8). Other enriched sequences include AT-rich simple motifs which are enriched in the GC-poor regions of the centromere islands [12] and heterochromatin in general [16]. Of particular note is the motif “TAGTTTT” which is both AT-rich and the reverse complement of the 5’ end of the

NTS fragment in *D. melanogaster*. This motif is frequently found on either end of *G2/Jockey-3* but not both ends simultaneously. The motif appears as an extension of the poly-A tail on the 3' end but appears to be a smaller part of the larger NTS fragment on 26 *G2/Jockey-3* insertions. These 5' NTS fragments were typically found on longer *G2/Jockey-3* elements suggesting they are typically part of the TE. Studies of human LINE-1 indicate transcription initiation for retroelements can vary considerably upstream and downstream of the promoter [16, 17]. Variable transcription initiation of *G2/Jockey-3* could incorporate a fragment of the surrounding NTS into subsequent insertions found elsewhere in the genome. However, this is unlikely because TAGTTTT was present in the reference genome prior to insertion for all but one of *de novo* insertions associated with the sequence (File S6). In addition, our phylogenetic analysis indicates that the *G2/Jockey-3* association with NTS does not have a single phylogenetic origin (Figure S2) and younger insertions were not more enriched for the NTS fragment than older fragments (FET, $p = 0.45$). This may be due to differences in insertion mechanisms; for instance, truncated L1 elements have different insertion preferences in comparison to full-length insertions [17, 18]. Surprisingly, the NTS sequences associated with *G2/Jockey-3* are distinct from the centromere-specific variants of NTS found at Cen3 in *D. melanogaster* [12].

*Gene conversion among Y-linked* G2/Jockey-3

Duplications of *G2/Jockey-3* and other intervening elements may be the products of gene conversion, ectopic recombination, or a combination of the two. Gene conversion leads to the homogenization of duplicated genes, repeats, and TEs [19, 20] and contributes to a pattern of concerted evolution [21]. On the *D. melanogaster* Y chromosome, recombination via crossing over is virtually absent and gene conversion occurs at higher rates than the autosomes [22, 23]. We estimated gene conversion between copies of *G2/Jockey-3* on the Y centromere, keeping in mind the classes and sizes of *G2/Jockey-3* fragments. Assuming a mutation rate of 2.8E-9 [24], we estimate gene conversion rates to range from 2.80E-5 to 7.54E-6 per site per generation among the different classes of Y-linked *G2/Jockey-3* insertions (Table S6). These rates of gene conversion are ~10E3 times higher than estimated gene conversion rates on the other chromosomes in *Drosophila* [25-27] and an order of magnitude less than previously estimated gene conversion rates between duplicated gene families on the Y chromosome [23]. Gene conversion rates are difficult to estimate, especially for TEs, because amplification via duplication or transposition and selection can mask true gene conversion events or be perceived falsely as conversion. There is also significant variation between *G2/Jockey-3* fragments within the same Y centromere classes suggesting homogenization of repeats via gene conversion is not complete (Figure S6).

### SUPPLEMENTAL FIGURES

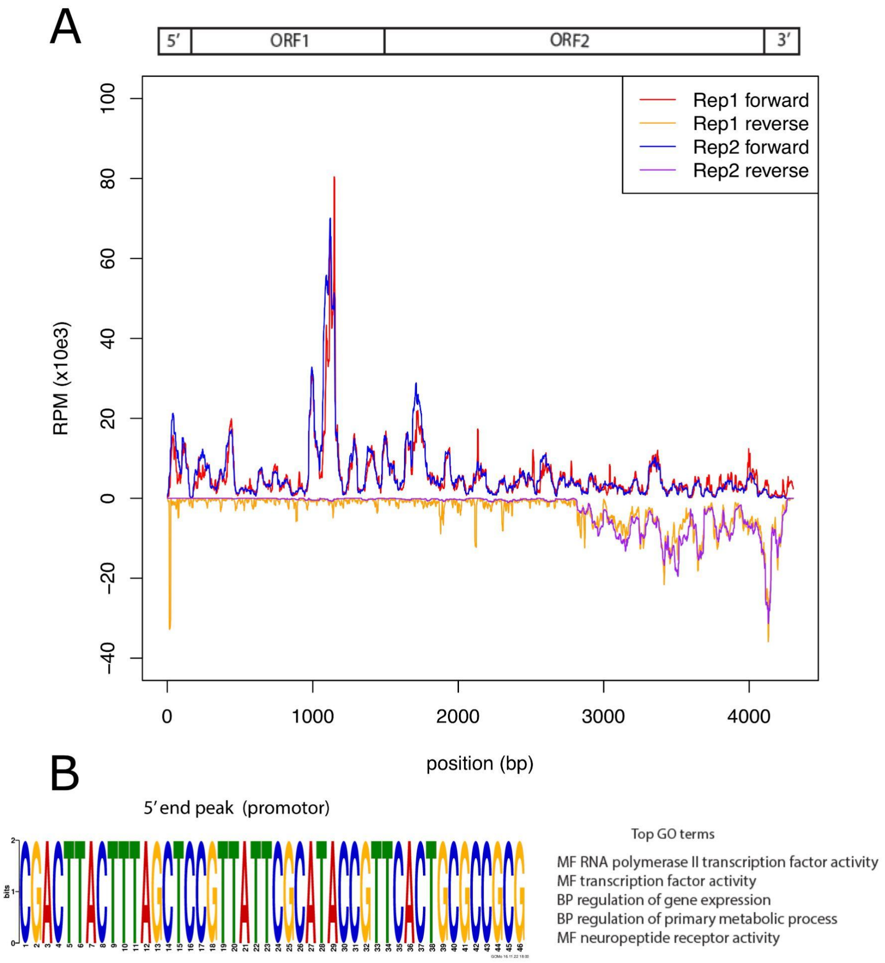


**Figure S1: Validation of the new annotation boundaries for *G2/Jockey3* using PROseq data and identification of a putative promoter at the 5' end of the element**. A) Normalized read depth (RPM) for PROseq data along the length of *G2/Jockey-3* with corresponding domains above. The line colors correspond to two replicate datasets (Rep1 and Rep2) and the orientation of the reads. ORF1 extends from 227-1519 bp and contains a zinc-binding domain. ORF2 overlaps slightly with ORF1 encompassing 1498-4197 bp and encodes the reverse transcriptase. B) GoMo analysis of the sequence in the 5' end of *G2/Jockey-3* corresponds to RNA polymerase II transcription activity and other transcription factor activity ($P < 0.01$).

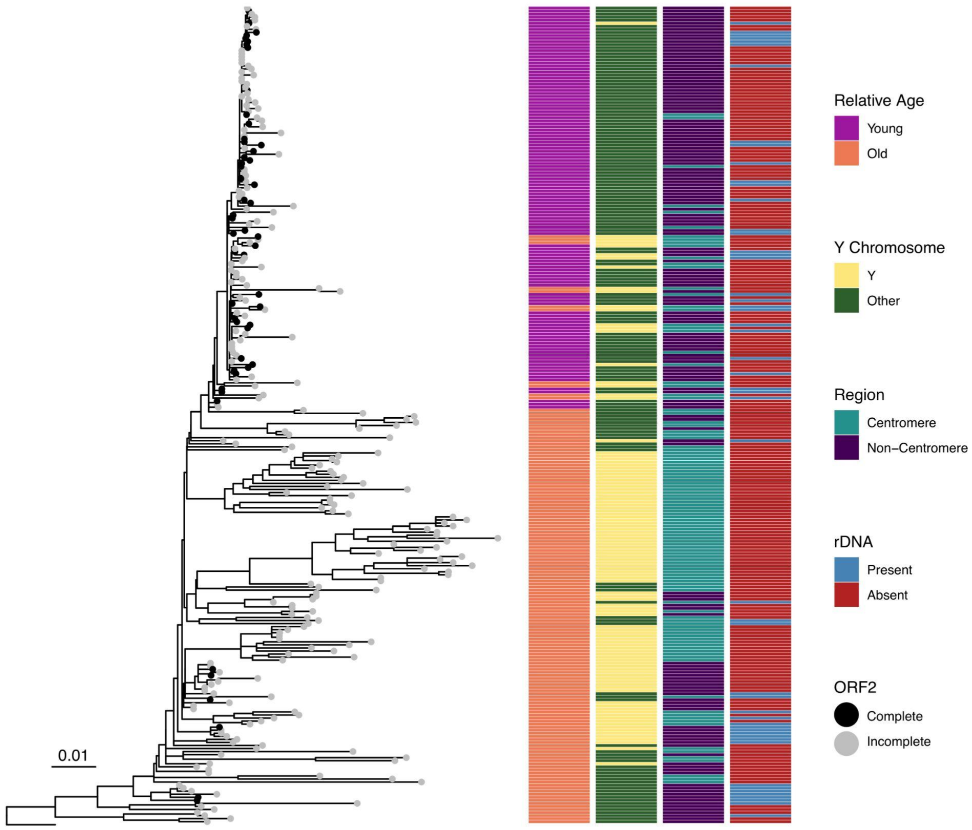


**Figure S2: Young *G2/Jockey-3* insertions share a common origin and are enriched in the centromeres; old and young insertions show a weak association with rDNA sequences.** A maximum likelihood phylogenetic tree with all non-redundant *G2/Jockey-3* sequences larger than 10% of the full-length (400 bp) in the reference genome and long-read samples with the *D. simulans* consensus as the outgroup. Black circles denote “complete” *G2/Jockey-3* elements containing a full ORF2; gray circles are incomplete elements. The columns and colors correspond to characteristics of each insertion including relative age, presence on Y chromosome, chromatin region, and presence of rDNA (both TAGTTTT and NTS sequences) within 15 bp of the insertion. Young insertions are those with < 1% divergence from the consensus sequence or < 2% and found within a single sample genome assembly.

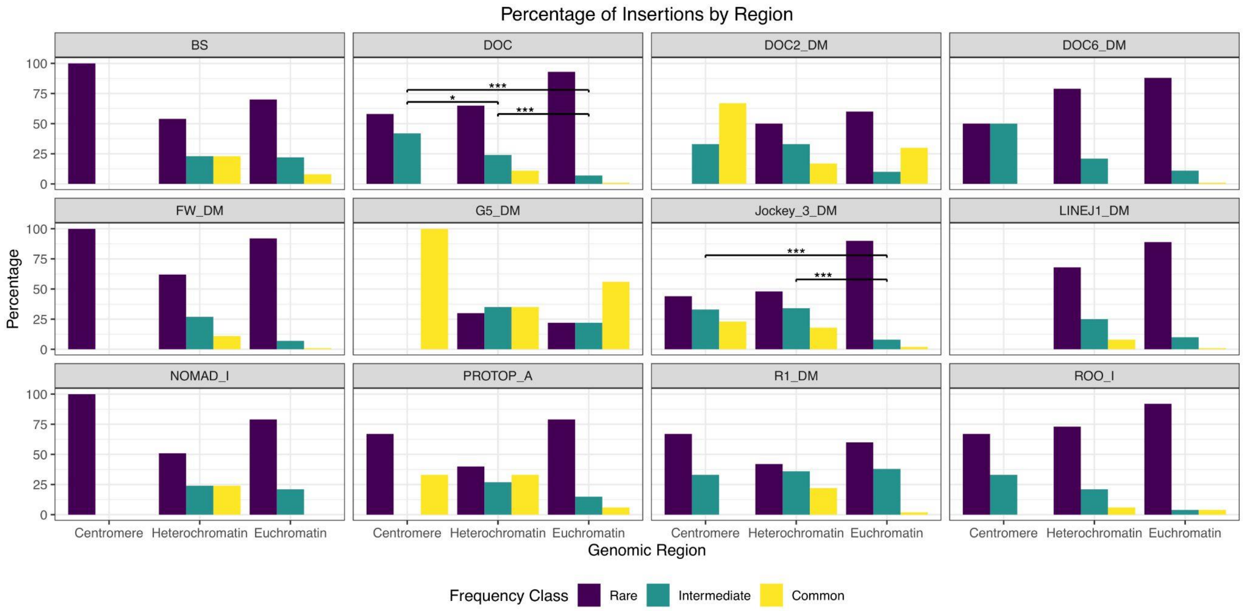


**Figure S3: Distribution of frequency variants for TE families with insertions in the centromere.** TE frequencies were binned into rare (less than 10% of all samples), intermediate (10-50% of all samples), and common (> 50% of all samples). Note that no statistics are reported where there are fewer than 5 insertions in any region. The frequency distributions for all of the TE families detected in the centromeres of the DGRP are compared between the centromere, heterochromatin, and euchromatin. Fisher's Exact Tests between regions, Bonferoni adjusted two-tailed P-value * $P < 0.05$, *** $P < 0.001$.

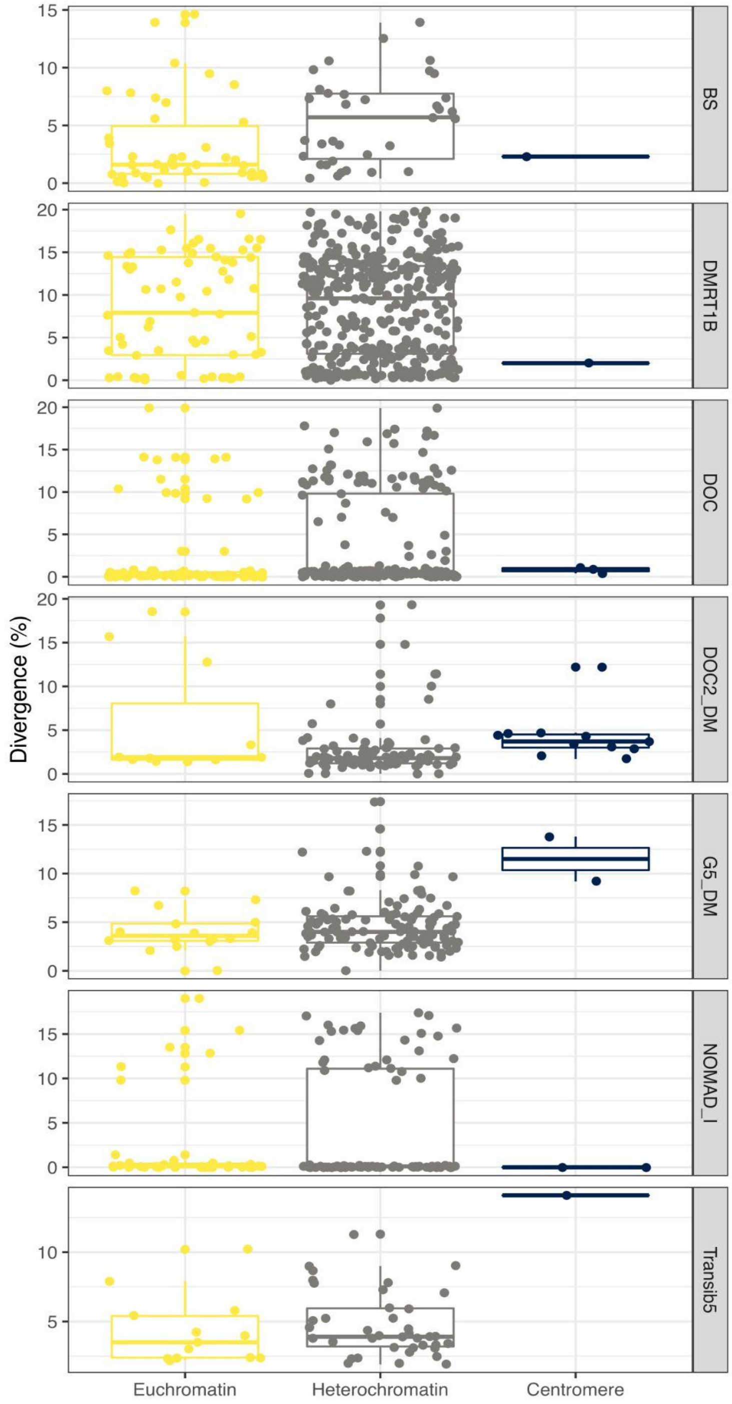


**Figure S4: Centromeric insertions in other TE families are older or align with increased activity of TE families in other heterochromatic regions.** The number of insertions for TEs other than *G2/Jockey-3* in the centromere grouped by divergence from the consensus sequence split between the euchromatin, heterochromatin and centromere.

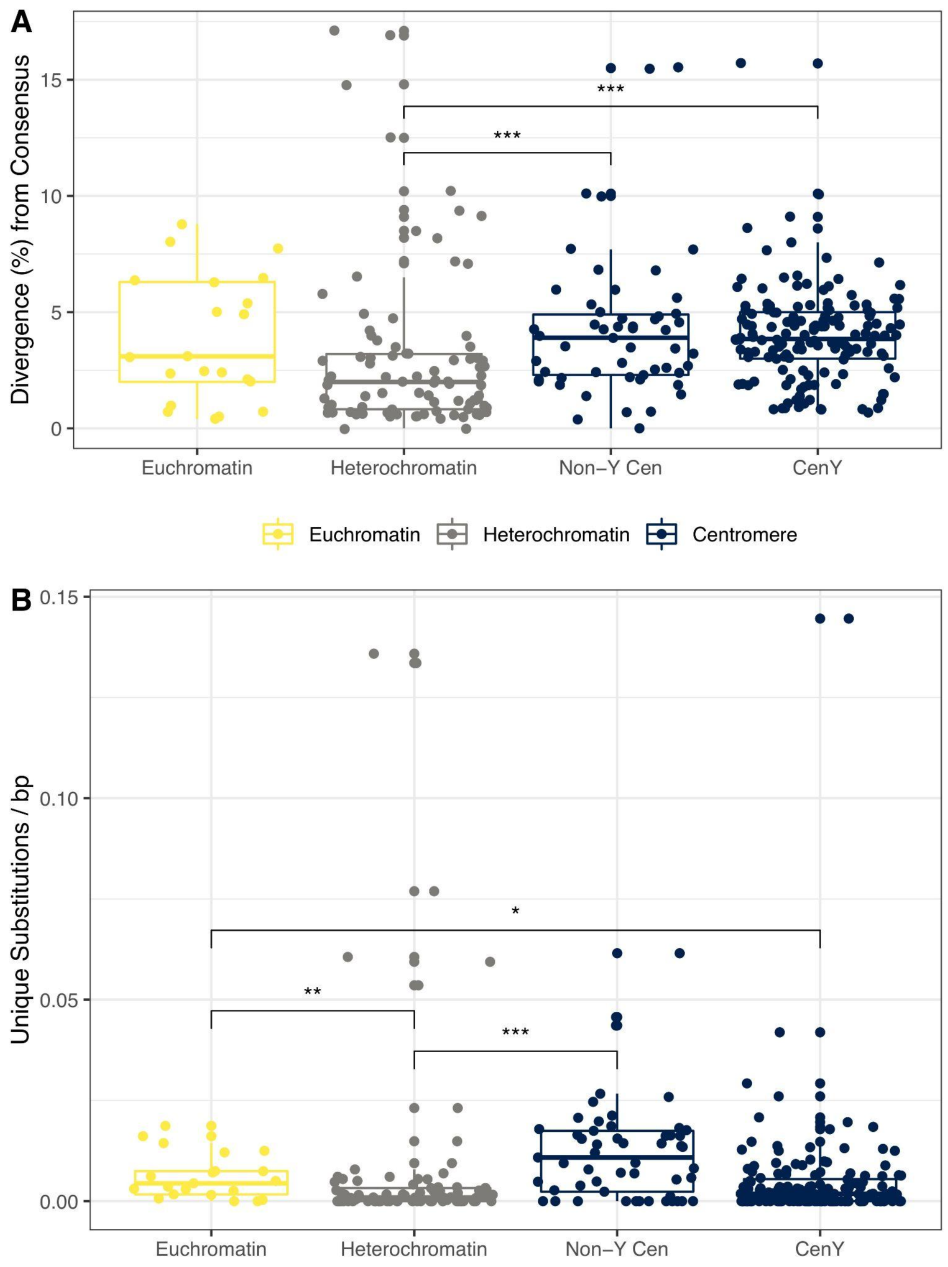


**Figure S5: The number of unique nucleotide substitutions in each *G2/Jockey-3* insertion divided by insertion length.** The points are colored by region; euchromatin is yellow, heterochromatin is gray, and centromere is navy. * $P < 0.05$, ** $P < 0.01$, *** $P < 0.001$. Non-significant comparisons are not shown.

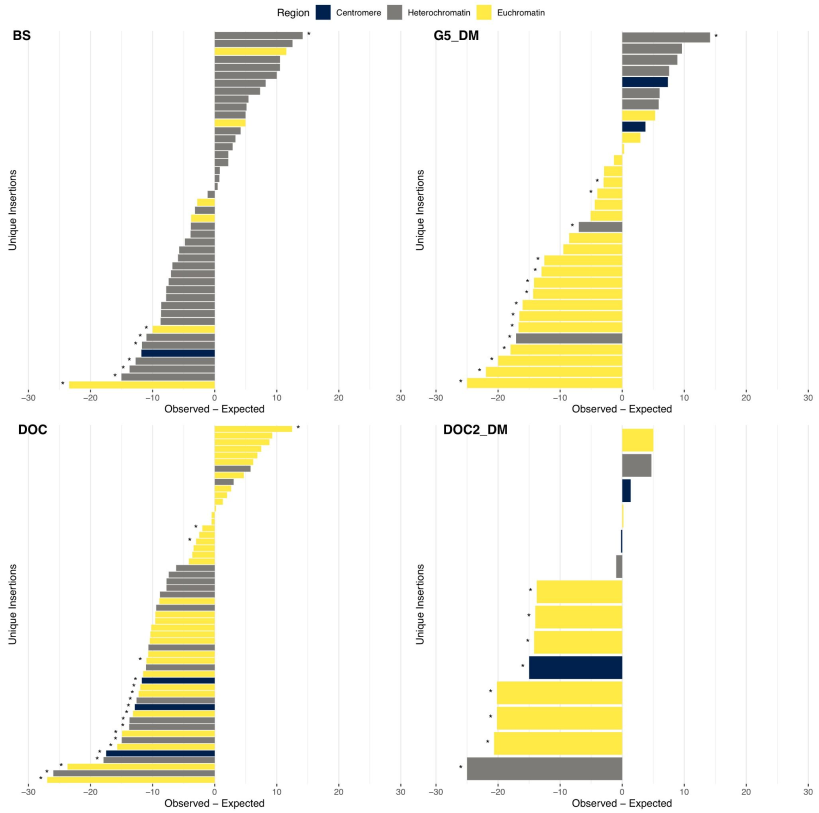


**Figure S6: Observed - expected frequencies of other TEs seen in the centromere islands suggest *G2/Jockey-3* shares the same selective pressure as other canonical, deleterious TE families.** The majority of TEs are found either less often than expected given age or are consistent with neutral expectations. The p-values are calculated as the sum of the probabilities that an insertion is found less frequent than expected (if observed - expected < 0) or more frequent than expected (observed - expected > 0). * $P < 0.1$, a stringent threshold given the reduced power to detect deviations from neutrality as stated in Blumenstiel et al. [28].

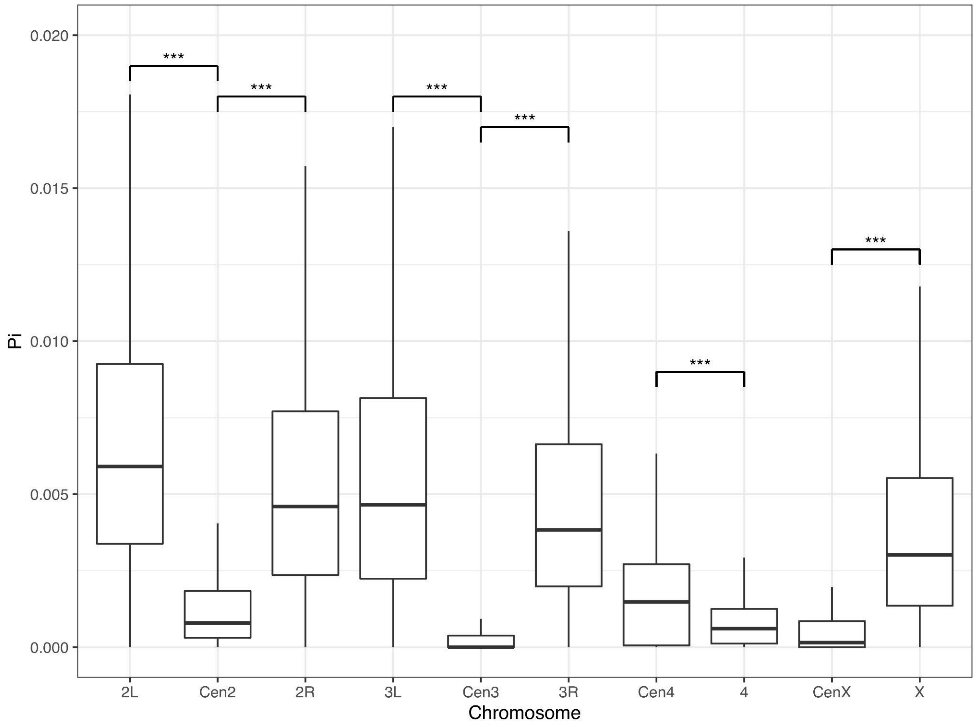


**Figure S7: Nucleotide diversity (pi) is significantly reduced in the centromere islands in comparison to the major chromosome arms.**

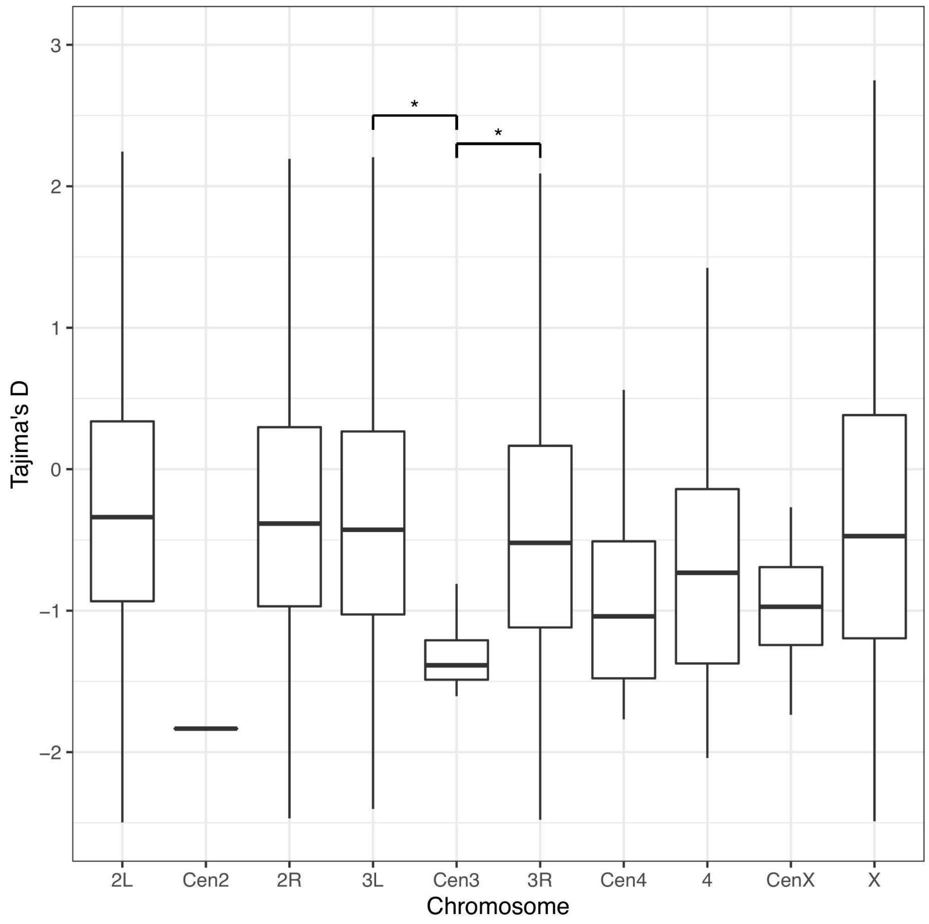


**Figure S8: Tajima's D is reduced in the centromere islands in comparison to the major chromosome arms except for Chromosome 4.** The fourth chromosome, like the centromere islands, experiences infrequent crossing over. Lower Tajima's D values are indicative of an abundance of rare alleles.

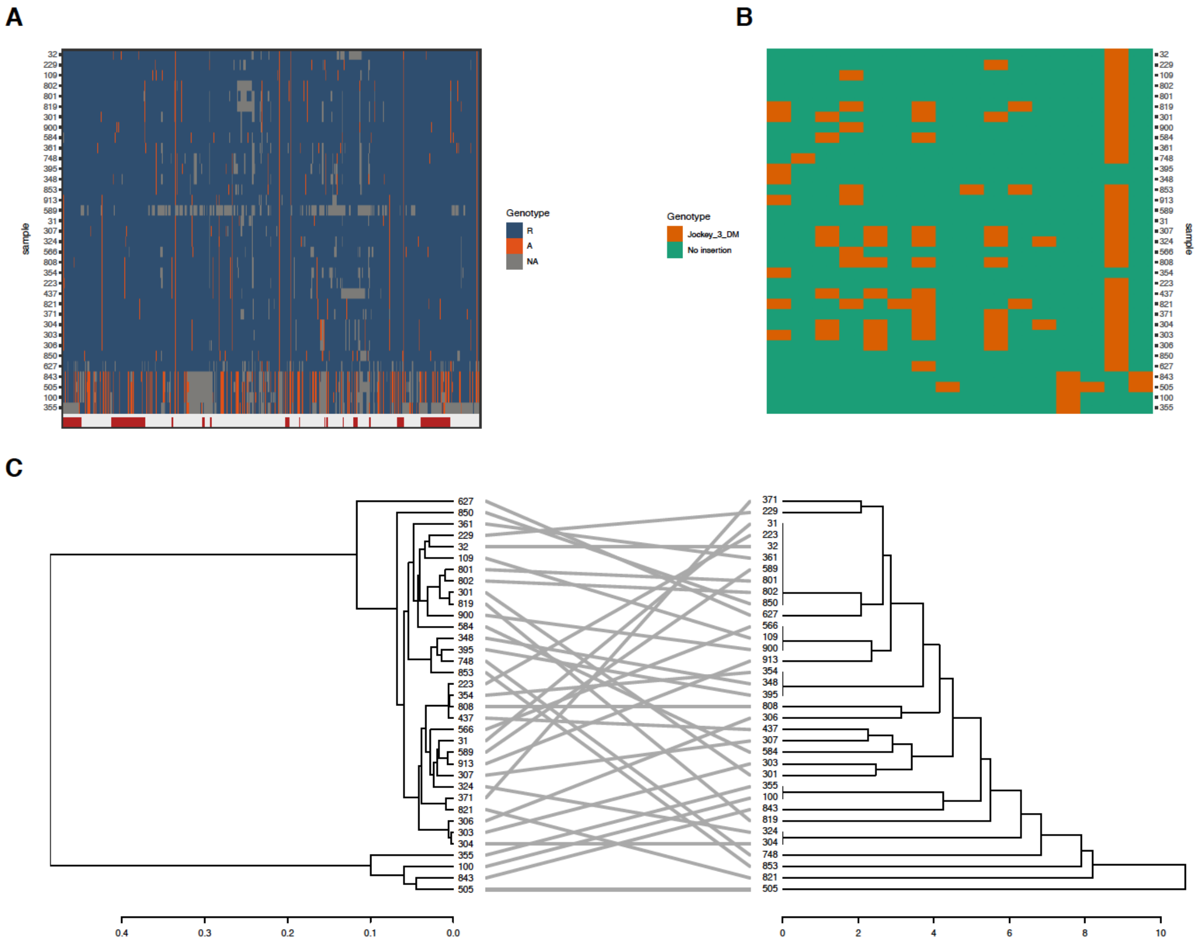


**Figure S9: Haplotypes do not correlate with *G2/Jockey-3* insertion polymorphism on the X chromosome centromere. A)** The colors correspond to variant sites along the length of the centromere islands in high coverage DGRP samples. The colors indicate whether the variant is the reference allele (dark blue), an alternative allele (red), or not available (gray). Annotation of *G2/Jockey-3* insertions from the reference genome (red) is shown below the haplotype blocks. **B)** The colors correspond to the presence (orange) and absence (green) of *G2/Jockey-3* insertions in high coverage DGRP samples along the length of CenX. **C)** Comparison of separate clustering analyses of SNP haplotypes on the left and *G2/Jockey-3* insertion polymorphisms on the right. Lines connect the samples from both analyses.

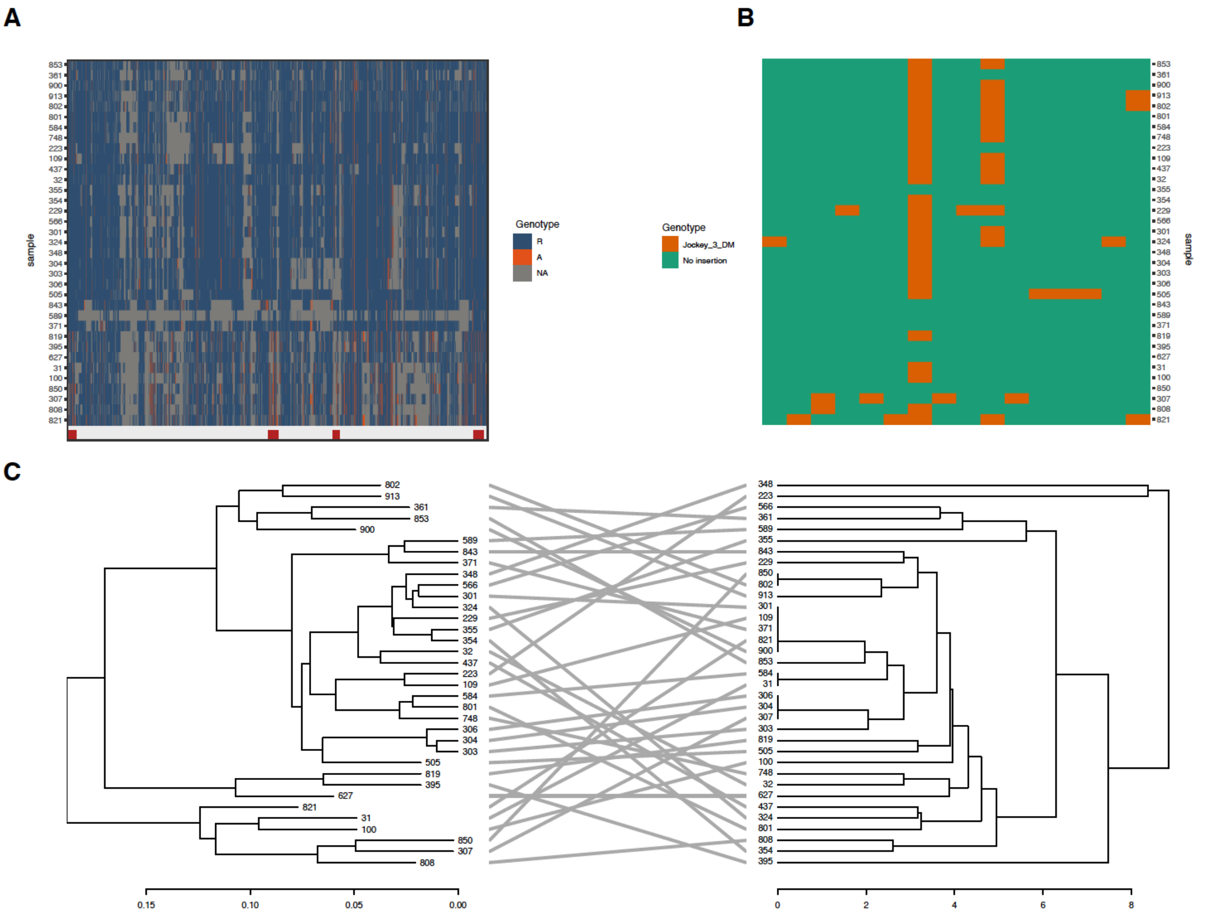


**Figure S10: Haplotypes do not correlate with *G2/Jockey-3* insertion polymorphism on the 3rd chromosome centromere. A)** The colors correspond to variant sites along the length of the centromere islands in high coverage DGRP samples. The colors indicate whether the variant is the reference allele (dark blue), an alternative allele (red), or not available (gray). Annotation of *G2/Jockey-3* insertions from the reference genome (red) is shown below the haplotype blocks. **B)** The colors correspond to the presence (orange) and absence (green) of *G2/Jockey-3* insertions in high coverage DGRP samples along the length of CenX. **C)** Comparison of separate clustering analyses of SNP haplotypes on the left and *G2/Jockey-3* insertion polymorphisms on the right. Lines connect the samples from both analyses.

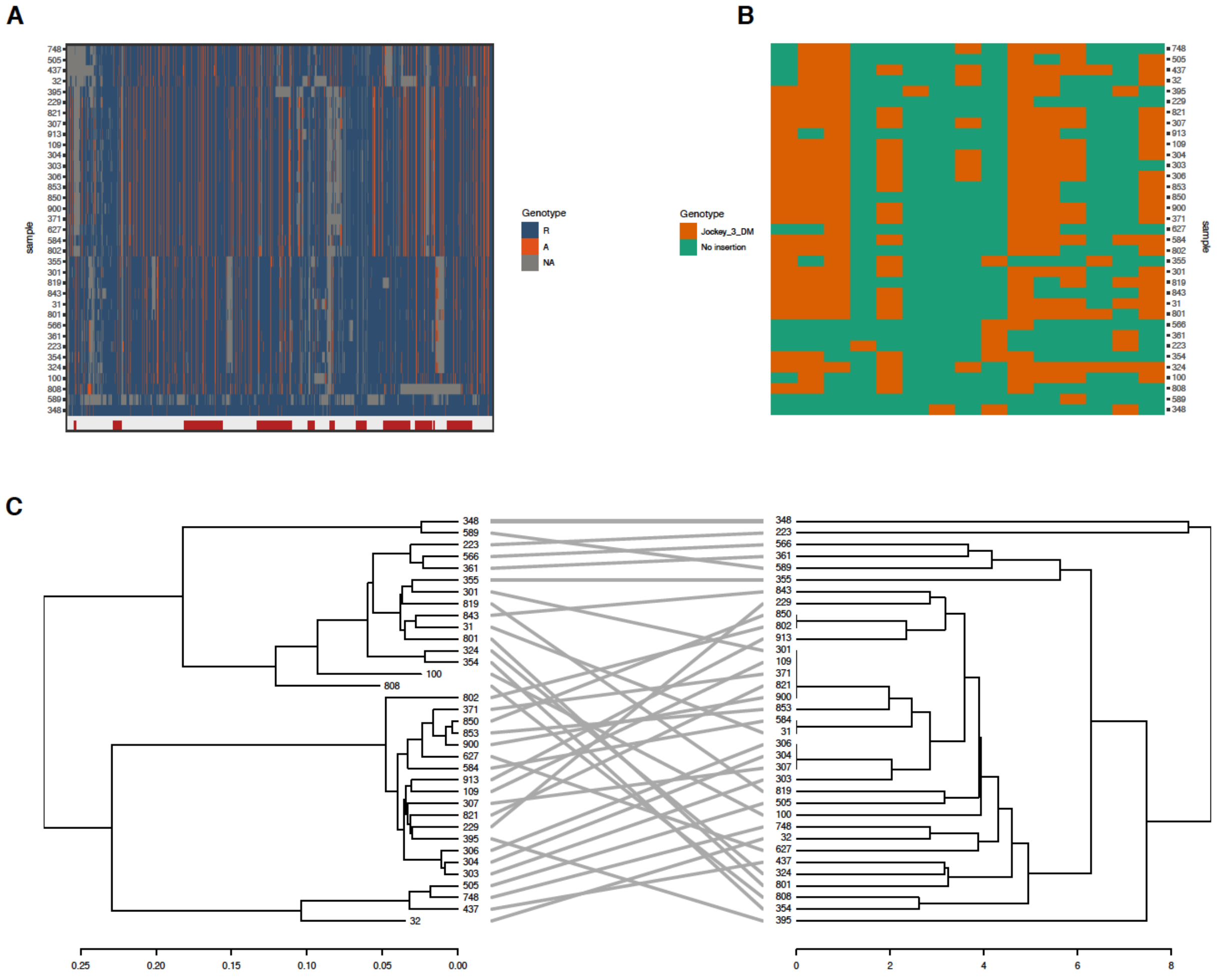


**Figure S11: Haplotypes do not correlate with *G2/Jockey-3* insertion polymorphism on the 4th chromosome centromere. A)** The colors correspond to variant sites along the length of the centromere islands in high coverage DGRP samples. The colors indicate whether the variant is the reference allele (dark blue), an alternative allele (red), or not available (gray). Annotation of *G2/Jockey-3* insertions from the reference genome (red) is shown below the haplotype blocks. **B)** The colors correspond to the presence (orange) and absence (green) of *G2/Jockey-3* insertions in high coverage DGRP samples along the length of CenX. **C)** Comparison of separate clustering analyses of SNP haplotypes on the left and *G2/Jockey-3* insertion polymorphisms on the right. Lines connect the samples from both analyses.

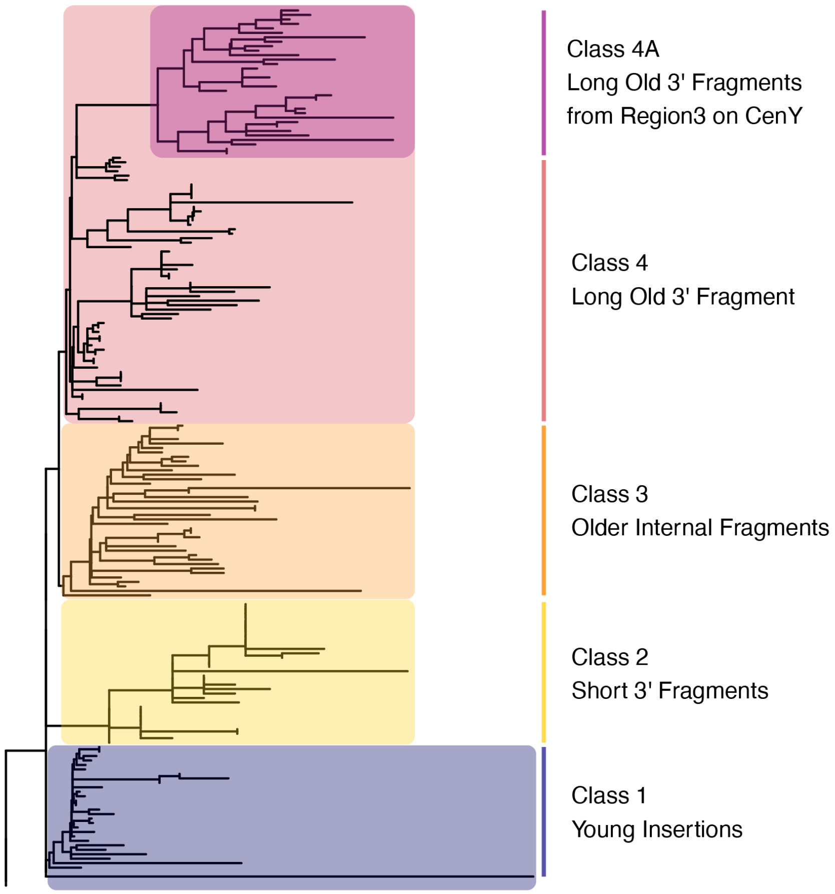


**Figure S12: Phylogenetic analysis groups all Y chromosome *G2/Jockey-3* sequences into four major clades and one subclade based on fragment size, divergence, and location.** We used these clades to define four classes of *G2/Jockey-3* elements on the Y chromosome: Class 1 younger long insertions, Class 2 short (< 160 bp) 3' end fragments, Class 3 older internal fragments missing the 5' and 3' terminal ends, Class 4 longer older 3' end fragments. Class 4 contains a subclade (4A) with highly similar sequences exclusively from the fourth region of CenY.

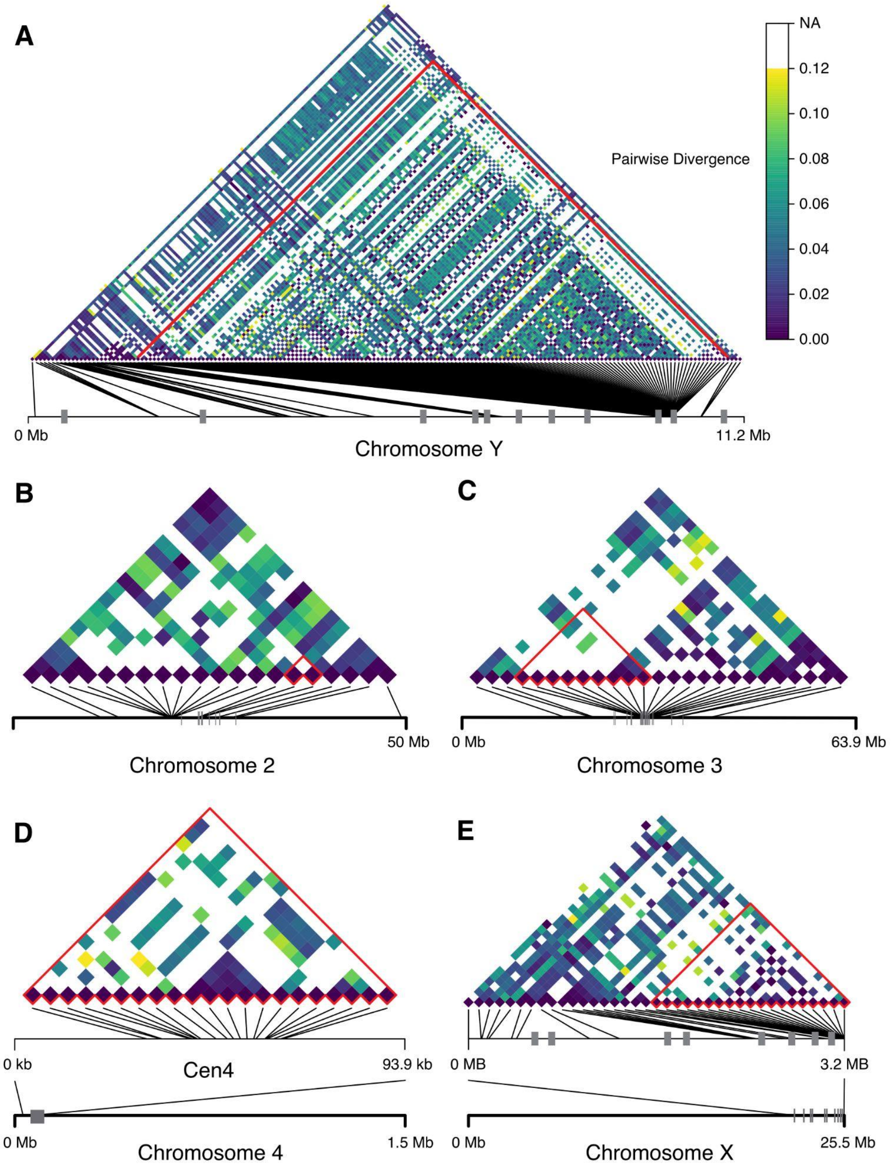


**Figure S13:** Pairwise divergence and relative positions of *G2/Jockey-3* along the length of every chromosome in *D. melanogaster*: A) Y Chromosome, B) Chromosome 2, C) Chromosome 3, D) Chromosome 4, and E) X Chromosome. Pairwise comparisons of centromeric copies of *G2/Jockey-3* are highlighted in red outline above. Cen4 and CenX with the surrounding regions are zoomed in as *G2/Jockey-3* is missing in most of the chromosome arms on the 4th and X chromosomes and to better understand the distribution of *G2/Jockey-3*. Gaps between contigs are displayed as gray boxes.

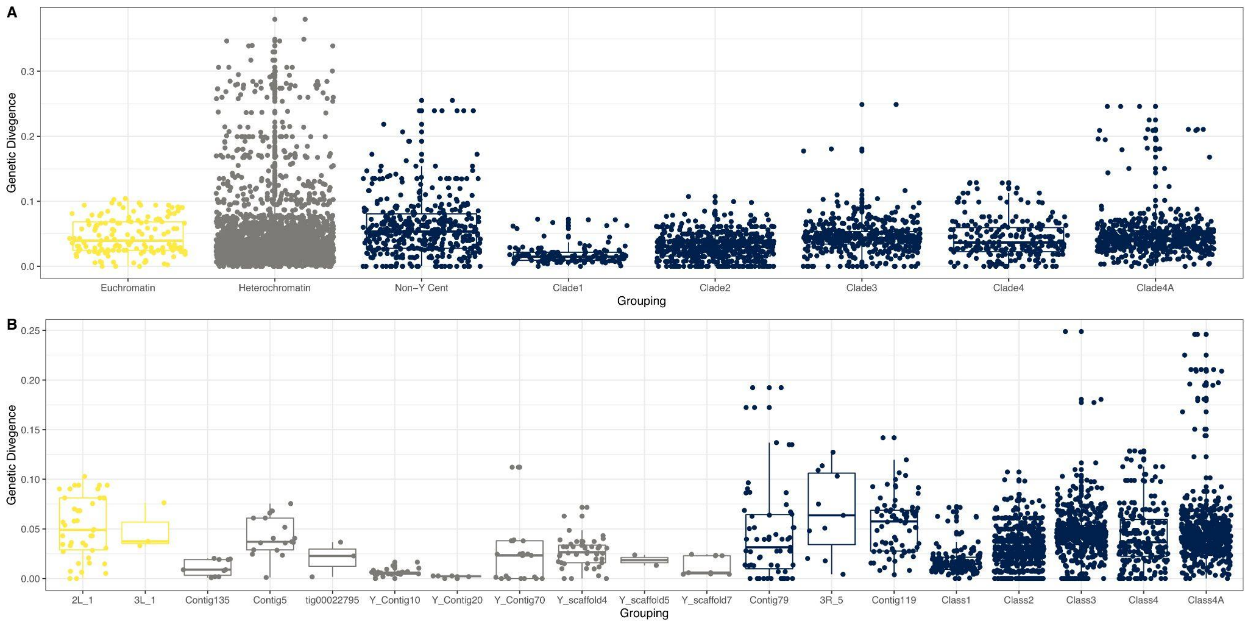


**Figure S14:** Pairwise divergence between copies of *G2/Jockey-3* A) divided into broad chromatin regions, non-Y centromeres, and broad classes within the Y centromere (based on the groupings from Figure S12). B) Pairwise divergences between *G2/Jockey-3* copies within the contigs. Color is based on chromatin, yellow contigs are majority euchromatin, gray is non-centromeric heterochromatin, and navy is centromeric.

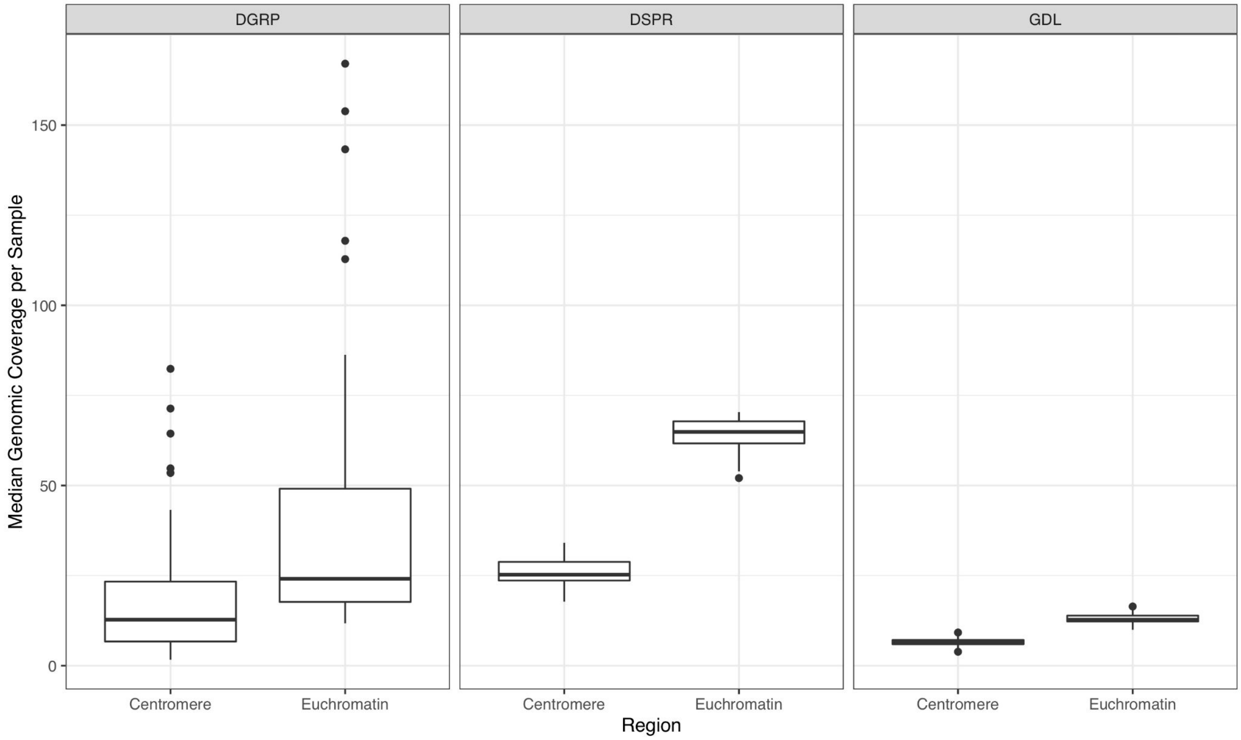


**Figure ST1: The median genomic coverage of the centromeres and euchromatin in samples from the DGRP, DSPR, and GDL.** Centromere coverage was consistently lower than the euchromatic regions of the genome. Samples in the GDL were also much lower than the DGRP and DSPR.

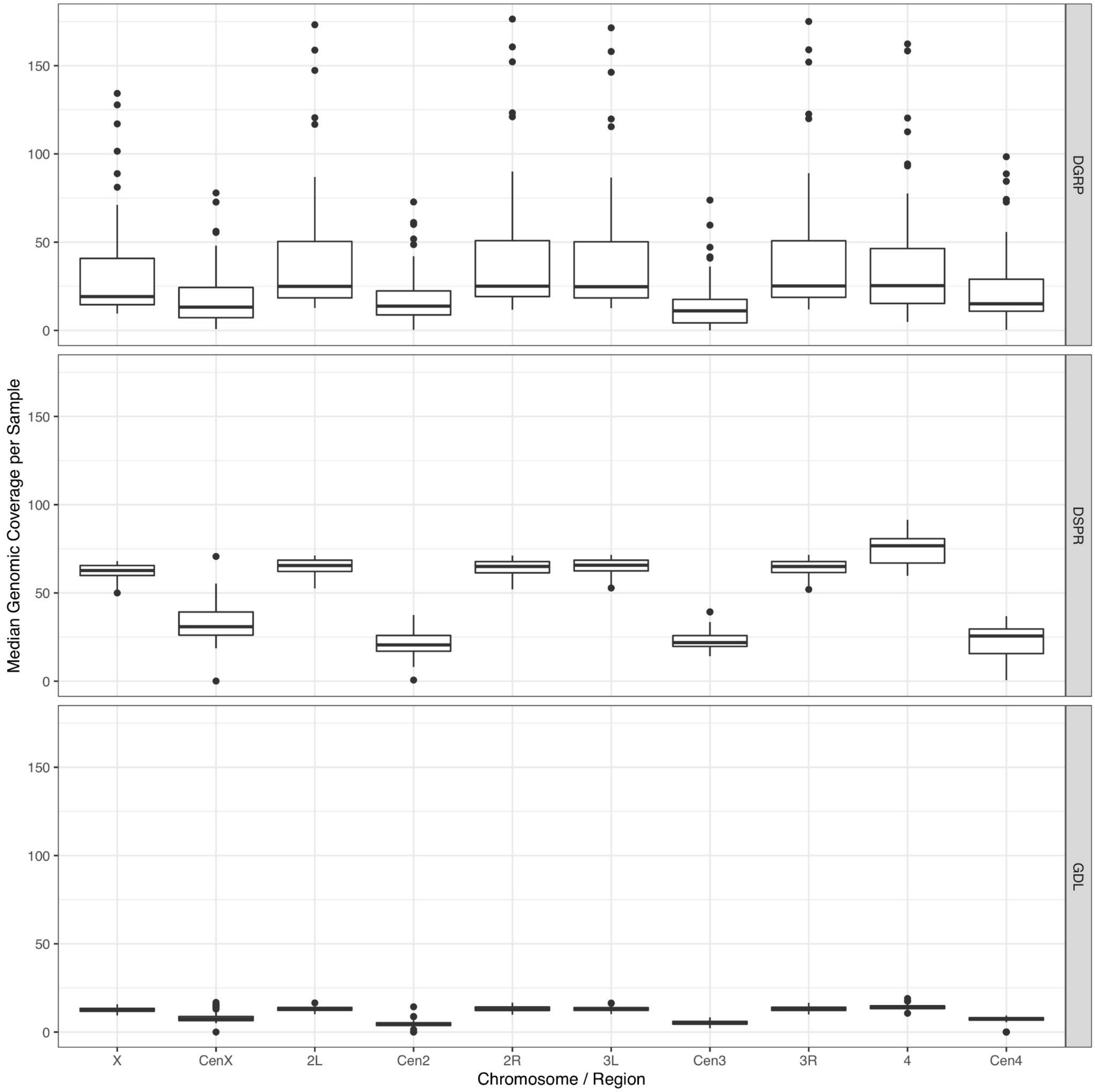


**Figure ST2: The median genomic coverage of the individual centromeres and chromosome arms in samples from the DGRP, DSPR, and GDL.** Centromere coverage was consistently lower than the euchromatic regions of the genome. Samples in the GDL were also much lower than the DGRP and DSPR. The X chromosome coverage also dips slightly in comparison to the autosomal arms.

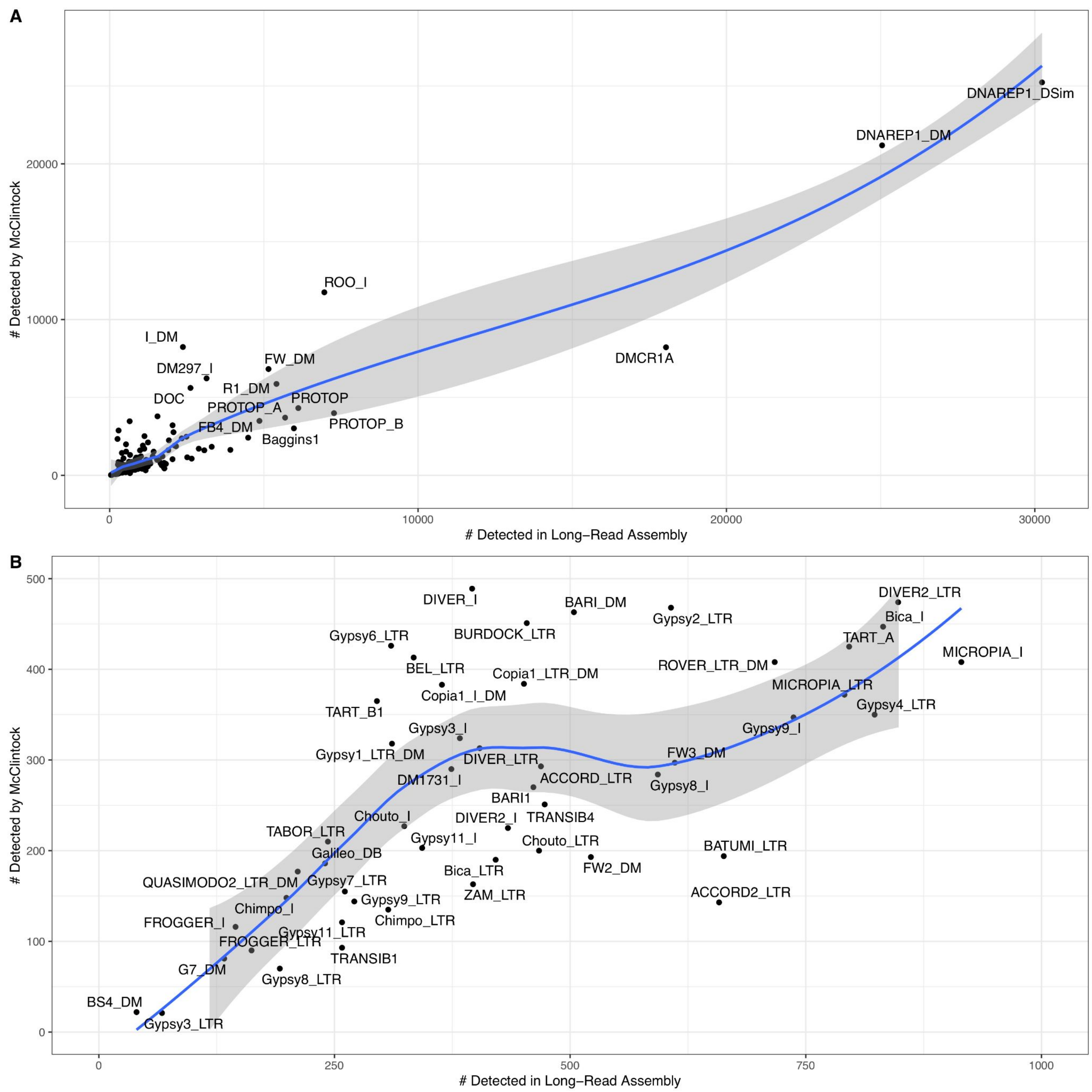


**Figure ST3: Long-read assemblies detected more reference TE insertions than the corresponding short-read data with the McClintock pipeline.** A) All TE families with some of the more common families labeled. B) The same graph zoomed in to visual the remainder of the TE families detected in both datasets.

**SUPPLEMENTAL TABLES**

**Table S1: Summary of data sources used in the study.**

**Table S2: Centromere sequences identified in long-read assemblies of *D. melanogaster* population samples.**

**Table S3: The average depth of short-read coverage in 500 bp windows for each long-read assembly line within the euchromatin and centromeres separately.**

**Table S4: RPM count of piRNAs mapped to TEs and satellite DNAs in *D. melanogaster* ovary and testis tissues.**

**Table S5: A summary of median and mean nucleotide diversity ($\pi$) and Tajima's D for the major chromosome arms and centromeres.**

**Table S6: Calculated gene conversion rates ($c_g$) for classes of *G2/Jockey-3* in the Y centromere.**

### SUPPLEMENTAL FILES

**File S1: Updated consensus sequence for *G2/Jockey-3* and its open reading frames (ORFs).**

**File S2: Table of *G2/Jockey-3* copies detected in short-reads generated assemblies of *D. melanogaster* lines in the DSPR.**

**File S3: Table of *G2/Jockey-3* copies detected in long-read generated assemblies of *D. melanogaster* lines in the DSPR and GDL.**

**File S4: TEs detected by the short-read TE-detection pipeline, McClintock, in the DGRP.**

**File S5: TEs detected by the short-read TE-detection pipeline, McClintock, in the DSPR.**

**File S6: Table of *G2/Jockey-3* insertions used for phylogenetic analysis.**

**File S7: Phylogenetic tree in NEWICK format of *G2/Jockey-3* insertions detected in long-read generated assemblies of *D. melanogaster*.**

**File S8: MEME motif analysis output of the 100 bp upstream and downstream of *G2/Jockey-3* insertions in the reference genome.**

**File S9: The combined input table used for the age-of-allele test and output of probabilities of observing a TE at a specific frequency in the population of high-coverage DGRP samples.**

**File S10: Drosophila TE family library for running the McClintock TE detection pipeline.**

**File S11: Table of *G2/Jockey-3* copies detected in five long-read generated assemblies of *D. melanogaster* lines in the DGRP.**

**File S12: Table of *G2/Jockey-3* copies detected in short-reads compared to the assemblies of *D. melanogaster* lines in the DGRP.**
